# Cleavage of *Streptococcus pneumoniae* ribosomal protein L27 by the Prp protease

**DOI:** 10.1101/2025.08.01.668195

**Authors:** Amarshi Mukherjee, Mohamed O. Nasef, Patrick M. Lindstrom, Vipin Chembilikandy, Carlos J. Orihuela, Terje Dokland

## Abstract

*Streptococcus pneumoniae* is one of the most important human respiratory pathogens worldwide. The increase in antibiotic resistance in *S. pneumoniae* and other pathogens is a significant public health concern. The streptococcal 70S ribosome is a prime target for antibiotics. Ribosomal protein L27 reaches into the peptidyl transferase center with its extended N-terminus and may be involved in the translation process. We have shown that L27 in Firmicutes, including staphylococci and streptococci, has an additional 9-12 amino acid N-terminal extension compared to Gram-negative organisms like *Escherichia coli*. The extension is cleaved by a protease called Prp that is absent from organisms that lack the extension. In *S. aureus*, Prp and the N-terminal extension of L27 are essential.

Here, we have characterized the cleavage of L27 by Prp in *S. pneumoniae*. Prp forms dimers that efficiently cleave L27 *in vitro*. An inactive form of Prp (PrpC34S) binds to L27 without cleaving, whereas L27 with a mutation (F12A) of the cleavage site does not bind Prp. Overexpression of PrpC34S *in vivo* is detrimental to *S. pneumoniae* growth. Surprisingly, a *S. pneumoniae* Δ*prp* strain was viable, apparently due to cleavage of L27 by another, unknown protease. Unlike in *S. aureus*, a mutant strain lacking the N-terminal extension of L27 was viable, but showed impaired growth. Our study sheds light on a process that could be exploited for novel antibiotics, but emphasizes important differences between streptococci and staphylococci.

**HIGHLIGHTS:**

- Ribosomal protein L27 in *S. pneumoniae* is N-terminally processed by Prp protease
- A *S. pneumoniae* Δ*prp* mutant is viable, but exhibits impaired growth
- Cleavage of L27 is required for viability, but the N-terminal extension is not essential
- In the absence of Prp, L27 is processed by another protease.
- There are distinct differences in the role of Prp between *S. aureus* and *S. pneumoniae*.

## INTRODUCTION

Members of the *Streptococcus* genus include many human pathogens of medical significance, including *S. pneumoniae, S. pyogenes, S. mutans*, and *S. agalactiae*. These pathogens can cause a wide range of conditions, including skin diseases, pneumonia, encephalitis, toxic shock and tooth decay. *S. pneumoniae* (pneumococcus) is one of the most important respiratory pathogens in humans [1–3], responsible for at least half a million deaths among children worldwide every year [4] and a common cause of bacterial meningitis [5]. The emergence of antibiotic resistance in *S. pneumoniae* and other bacteria is a significant concern [6–8]. Development of new antibiotics has not kept up with the increase in resistance. Furthermore, streptococci can be induced to a reversible persister phenotype that is tolerant to antibiotics [9, 10]. The mechanisms underlying such tolerance are poorly understood.

The bacterial ribosome is a prime target for numerous antibiotics, including tetracyclines, chloramphenicol, lincosamides (e.g. clindamycin), macrolides (e.g. erythromycin), aminoglycosides (e.g. streptomycin) and oxazolidinones (linezolid) [11–14], many of which are in use against *S. pneumoniae* and other Gram-positive pathogens [15]. The bacterial ribosome (70S) consists of a large (50S) and a small (30S) subunit, and is comprised of 56 proteins labeled L1–L36 and S1–S22 and three RNAs [16]. The catalytic steps in the translation process are carried out primarily by the RNA; however, the protein components are necessary for correct folding and assembly of the ribosome and may play roles in modulating its activity [16, 17]. L27, product of the *rpmA* gene, is a ribosomal large subunit protein that does not have an equivalent in eukaryotes and archaea. L27 is uniquely positioned in the ribosome where its N-terminus reaches into the peptidyl transferase center and interacts with the tRNA [18]. However, its involvement in peptide bond formation or other aspects of the translation process is still unclear [19–21].

In the Firmicutes, including streptococci as well as staphylococci, listeriae, bacilli, and clostridia, L27 has an additional, conserved 9-12-residue N-terminal extension that is not present in L27 from Gram-negative organisms like *Escherichia coli* [22, 23](Fig. 1A). We previously described a novel cysteine protease that cleaves the N-terminal extension of L27 in *Staphylococcus aureus* [23]. This protease is encoded by a gene that is located between *rplU* (encoding L21) and *rpmA* in the same operon. Because we originally identified this protein for its role in cleavage of bacteriophage proteins, we called it Prp, for “<u>P</u>hage-related ribosomal <u>p</u>rotease” [23]. The *prp* gene is present in the genomes of all Firmicutes and other bacteria that encode L27 with the N-terminal extension, but not in *E. coli* and other bacteria that lack the N-terminal extension [23]. Both the presence of the N-terminal extension of L27 and its ability to be cleaved were found to be essential in *S. aureus* using an expression system in which the wildtype L27 could be turned off and replaced with either truncated or uncleavable L27 [24]. Furthermore, overexpression of a protease-deficient mutant form of Prp (C34A) in *S. aureus* was inhibitive to growth and led to accumulation of ribosomal precursors [23], suggesting a defect in ribosome assembly.

**Figure 1.**
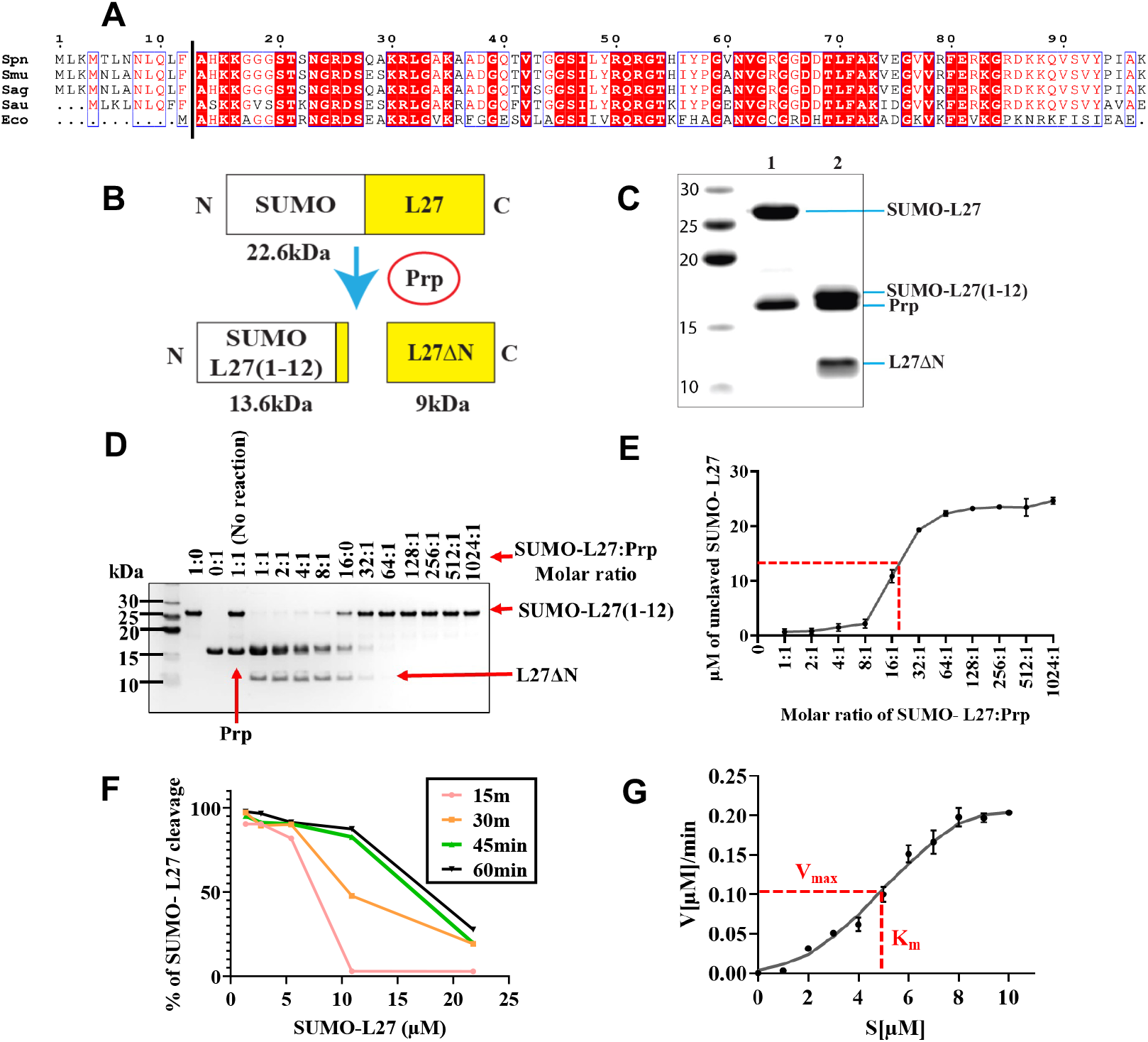
In vitro cleavage of L27. **(A)** Aligned amino acid sequences of L27 proteins from *S. pneumoniae* (Spn), *S. mutans* (Smu), *S. agalactiae* (Sag), *S. aureus* (Sau), and *E. coli* (Eco). The F|A cleavage site is shown as a vertical line. (**B**) Schematic representation of the in vitro cleavage of SUMO-L27 by Prp. (**C**). Analysis of SUMO-L27 digestion by Prp using 15% SDS-PAGE. Equimolar amounts of SUMO-L27 and Prp were mixed, immediately boiled in Laemmli sample buffer, and loaded onto SDS-PAGE (lane 1). An identical sample was incubated in 25 mM HEPES buffer (pH 7.4) containing 0.3 M NaCl at 37 °C for one hour before loading (lane 2). Lane M is marker; molecular weights (kDa) as indicated. **D**). Dose-dependent cleavage of SUMO-L27 by Prp. SUMO-L27 (22 µM) was incubated with increasing concentrations of Prp at 37 °C for one hour, and the reaction products were analyzed by 15% SDS-PAGE. (**E**). Graphical representation of the dose-dependent cleavage of SUMO-L27 by Prp, calculated from the gel. Concentration of uncleaved SUMO-L27 was plotted as a function of SUMO-L27:Prp ratio. The halfway point of the curve, corresponding to a ratio of 1:20 is indicated by the red dotted lines. (**F**) Time-dependent activity of Prp with varying concentrations of SUMO-L27, expressed as the percentage cleaved SUMO-L27 measured by SDS-PAGE. (**G**). Michaelis-Menten plot of the rate of SUMO-L27 cleavage (in µM/min) as a function of SUMO-L27 concentration (µM) in the presence of 1 µM Prp. The halfway point corresponding to K_m_=4.7 µM and V_max_ = 0.10 µM/min is indicated by the red lines.

Here, we show that Prp cleaves L27 *in vivo* and *in vitro* in *S. pneumoniae*. Surprisingly, *prp* is not essential in *S. pneumoniae*, although a Δ*prp* mutant exhibits impaired growth. This appears to be due to inefficient cleavage of L27 by some other cellular protease. Expression of an inactive form of Prp (PrpC34S) was detrimental to growth, apparently due to binding to L27 without cleavage and release. A mutant strain lacking the N-terminal extension of L27 was viable, showing that while cleavage of the N-terminal extension is essential, the extension itself is not required for viability, and indicating that there are distinct differences in the role of Prp and L27 between *S. aureus* and *S. pneumoniae*.

## RESULTS

### 1. Cleavage of L27 by Prp in vitro

To analyze the *in vitro* activity of Prp and its cleavage of L27, we first produced *S. pneumoniae* strain TIGR4 Prp with a C-terminal myc-His_6_ tag by expression in *E. coli*. We refer to this protein as Prp. We also generated a plasmid expressing full-length *S. pneumoniae* L27 with an N-terminally His_6_-tagged SUMO domain fused at its N-terminus, producing protein SUMO-L27. Both proteins were purified to homogeneity by affinity and size exclusion chromatography (SEC).

Cleavage of the 22.6 kDa SUMO-L27 fusion protein at the expected Prp cleavage site predicted by sequence comparison to *S. aureus* (MNLQFF|A) (Fig. 1A) would lead to two bands at 13.6 and 9.0 kDa, corresponding to the SUMO moiety with the first 12 amino acids of L27 [SUMO-L27(1-12)], and the mature L27 protein (L27ΔN; residues 13–97), respectively (Fig. 1B). When the SUMO-L27 protein was mixed with an equimolar amount of Prp, it was fully cleaved after 1 hr at 37 °C (Fig. 1C). The cleavage site was confirmed by full-length mass spectrometry to correspond to the expected cleavage site between Phe 12 and Ala 13 (Fig. S1).

SUMO-L27 (22 µM) was mixed with Prp at molar ratios ranging from 1:1 to 1024:1 (SUMO-L27:Prp), incubated at 37 °C for 1 hour in assay buffer (25 mM HEPES, pH 7.4, 0.3 M NaCl), and analyzed by SDS-PAGE (Fig. 1D). The results indicate that 1 µM Prp cleaves approximately 50% of SUMO-L27 within 1 hour under these conditions (Fig. 1E). We also carried out a time course experiment, in which Prp at 1.36 µM concentration was mixed with SUMO-L27 at molar ratios from 1:1 to 1:20 and incubated at 37°C for 15–60 mins (Fig. S2). Significant cleavage was observed within 15 minutes and continued for at least 60 minutes (Fig. 1F, S2).

To determine the Michaelis constant (K_M_), 1 µM Prp was incubated with varying concentrations of SUMO-L27 for 15 minutes at 37 °C and separated by SDS-PAGE (Fig. S3). The rate of cleavage was calculated (µM/min) and plotted against the SUMO-L27 concentration (Fig.1G). From this data we calculated a K_M_=5.0±0.2 µM, V_max_=0.1±0.1 µM/min and k_cat_=0.0017±0.0010 s^-1^, comparable to measurements on *S. aureus* Prp using a peptide substrate [25].

### 2. PrpC34S binds to L27 without releasing

The overexpressed SUMO-L27 and Prp proteins were separated by SEC on a Superdex™ 75 column. Prp and SUMO-L27 eluted as single peaks in fraction 6, consistent with the expected Prp dimer (33.4 kDa) [25, 26] and a likely SUMO-L27 monomer (22.6 kDa)(Fig. 2A). When the proteins were mixed, two new peaks appeared in fractions 7 and 8 (Fig. 2A). The peaks were confirmed by SDS-PAGE to correspond to the cleaved SUMO-L27 fragments, SUMO-L27(1-12) and L27ΔN, respectively (Fig. 2A), showing that the SUMO-L27 protein had already been fully cleaved by Prp prior to separation, and that Prp does not remain bound to L27 after cleavage.

**Figure 2.**
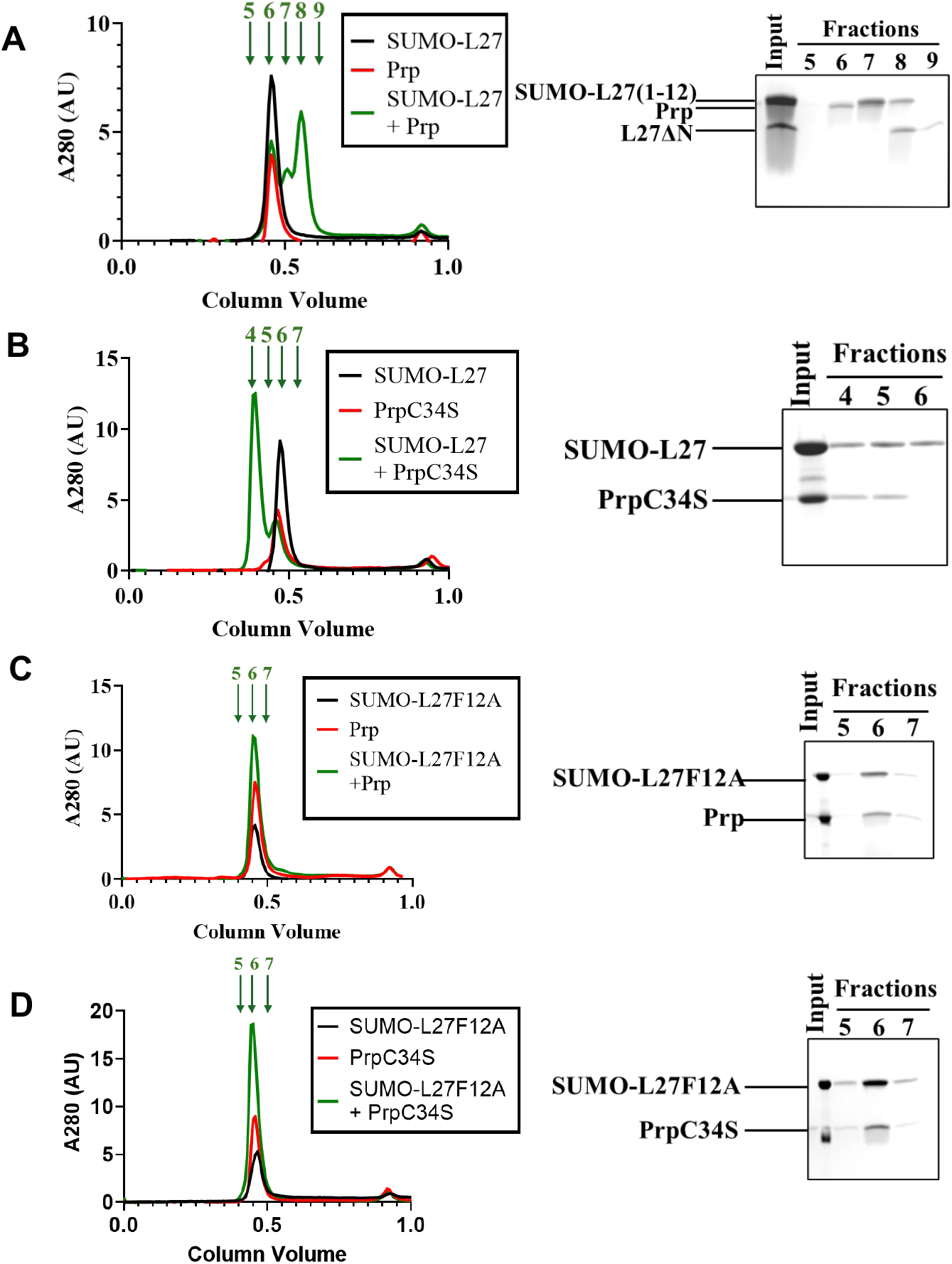
Separation of wildtype and mutant SUMO-L27 and Prp mixtures by size exclusion chromatography (SEC). **(A)** SUMO-L27 was mixed with wildtype Prp, and incubated at 37°C for 2 hrs, followed by separation by SEC (green curve). SUMO-L27 and Prp alone (black and red curves, respectively) were run as internal controls. The curve shows the appearance of two new peaks in fraction 7 and 8. SDS-PAGE of fractions 5-9, confirming that the fractions 7 and 8 contained the cleaved SUMO-L27 fragments, SUMO-L27(1-12) and L27ΔN. **(B)**. A mixture of SUMO-L27 with PrpC34S was prepared and separated in the same way as in (A). A new peak appeared in fraction 4. SDS-PAGE of fractions 4-6 confirmed that this peak contained uncleaved SUMO-L27 and PrpC34S, showing that the two proteins bound without cleavage. **(C)** SEC separation of a mixture of the SUMO-L27F12A mutant and wildtype Prp. No new peaks appeared, indicating that there was no cleavage or interaction of the two proteins. Fractions 5-7 were separated by SDS-PAGE, showing the presence of uncleaved SUMO-L27F12A and Prp in fraction 6. **(D)** SEC separation of SUMO-L27F12A mixed with PrpC34S, showing no cleavage and no interaction.

Previous results from *S. aureus* had shown that Prp with mutation of the active site Cys residue (PrpC34A and PrpC34S) was inactive [23, 25]. We generated plasmids overexpressing C-terminally myc-His_6_-tagged *S. pneumoniae* Prp with mutations of the predicted active site Cys 34 to either Ala (PrpC34A) or Ser (PrpC34S). Both proteins were purified the same way as Prp and likewise eluted as dimers by SEC. When PrpC34S was mixed with SUMO-L27, no cleavage occurred, even after 2 hrs of incubation (Fig. S4A,B). Like *S. aureus* Prp [25], *S. pneumoniae* Prp was not inhibited by the cysteine protease inhibitor E-64 (Fig.S4A).

The mixture of SUMO-L27 and PrpC34S (at 1:1 molar ratio) was then separated by SEC. A new major peak appeared in fraction 4, which was shown by SDS-PAGE to contain PrpC34S and uncleaved SUMO-L27 (Fig. 2B). The position of the peak was consistent with a complex consisting of a PrpC34S dimer with either one or two copies of SUMO-L27 (56.0 kDa and 78.6 kDa, respectively). Some excess SUMO-L27 remained unbound in fraction 6. This result showed that the mutant PrpC34S protein remained bound to SUMO-L27 without cleaving (Fig. 2B).

We also generated a SUMO-L27 construct with a Phe 12 to Ala mutation (F12A) in the predicted F|A cleavage site (SUMO-L27F12A). When this protein was mixed with either Prp or PrpC34S, peaks corresponding to neither the SUMO-L27 cleavage products nor the SUMO-L27:Prp complex appeared, showing that SUMO-L27F12A was unable to bind to either Prp or PrpC34S and did not become cleaved by Prp (Fig. 2C, D).

### 3. Deletion of prp causes a growth defect in S. pneumoniae

To investigate the role of Prp *in vivo*, we used homologous recombination to produce a *prp* knockout strain (Δ*prp*) in *S. pneumoniae* TIGR4, where *prp* (TIGR4 gene SP_1106) was replaced with a kanamycin resistance cassette (gene *aphIII*), producing the kanamycin resistant (Kan^R^) strain MN001 (Table 1, Figure 3). Whole-genome sequencing confirmed that the *prp* gene had been replaced by *aphIII* in its original locus (Fig. 3). This mutant was viable, but showed strongly delayed growth (Fig. 4A), with a twofold increase of the lag phase time compared to the TIGR4 wildtype (Table 2). During exponential growth, the Δ*prp* mutant had a similar generation time (63 mins) to the wildtype (61 mins); however, the Δ*prp* mutant never reached the same cell density as TIGR4 (Fig. 4A).

**Table 1.** Strains and plasmids used in this study.

| Strain name | Background | Genotype | Phenotype¶ | Reference |
| --- | --- | --- | --- | --- |
| TIGR4 |  | Virulent clinical capsular serotype 4 isolate. | wildtype | [39] |
| TIGR4S | TIGR4 | Mutation in the *rpsL* gene (ribosomal protein S12). | *rpsL*<sup>+</sup>, Sm<sup>R</sup> | [28] |
| MN001 | TIGR4 | *prp::kan<sup>R</sup>* | $\Delta prp$; Kan<sup>R</sup> | This study |
| MN009 | TIGR4S | *spv_2417::Spec<sup>R</sup>-P<sub>lac</sub>-prp<sup>WT</sup>-myc-his*;<br>*prp::kan<sup>R</sup>*<br>*prsA::Gm-PF6-lacI*; | LacI<sup>+</sup>, IPTG-inducible Prp-myc-his in a $\Delta prp$ background.<br>Sm<sup>R</sup>, Spec<sup>R</sup>, Kan<sup>R</sup>, Gm<sup>R</sup> | This study |
| MN011 | TIGR4S | *prsA::Gm<sup>R</sup>-PF6-lacI*;<br>*spv_2417::Spec<sup>R</sup>-P<sub>lac</sub>-prp<sup>C34S</sup>-myc-his* | IPTG-inducible expression of PrpC34S (myc-his) in a TIGR4S (Prp<sup>WT</sup>) background. Sm<sup>R</sup>, Gm<sup>R</sup>, Spec<sup>R</sup> | This study |
| Plasmid name |  |  |  |  |
| pPEPY-PF6-lacI |  |  | LacI under constitutive PF6 promoter; Gm<sup>R</sup>, Kan<sup>R</sup> | [42]. |
| pPEPZ-Plac |  |  | Used to construct IPTG-inducible *prp* strains. | [27] |
| pPEPZ-Plac-prp |  |  | Prp<sup>WT</sup> under lac promoter; Spec<sup>R</sup>; used to construct MN009 | This study |
| pPEPZ-Plac-prp |  |  | PrpC34S under lac promoter; Spec<sup>R</sup>; used to construct MN009 | This study |

¶Sm<sup>R</sup>, streptomycin resistance; Kan<sup>R</sup>, kanamycin resistance; Gm<sup>R</sup>, gentamycin resistance; Spec<sup>R</sup>, spectinomycin resistance.

**Figure 3.**
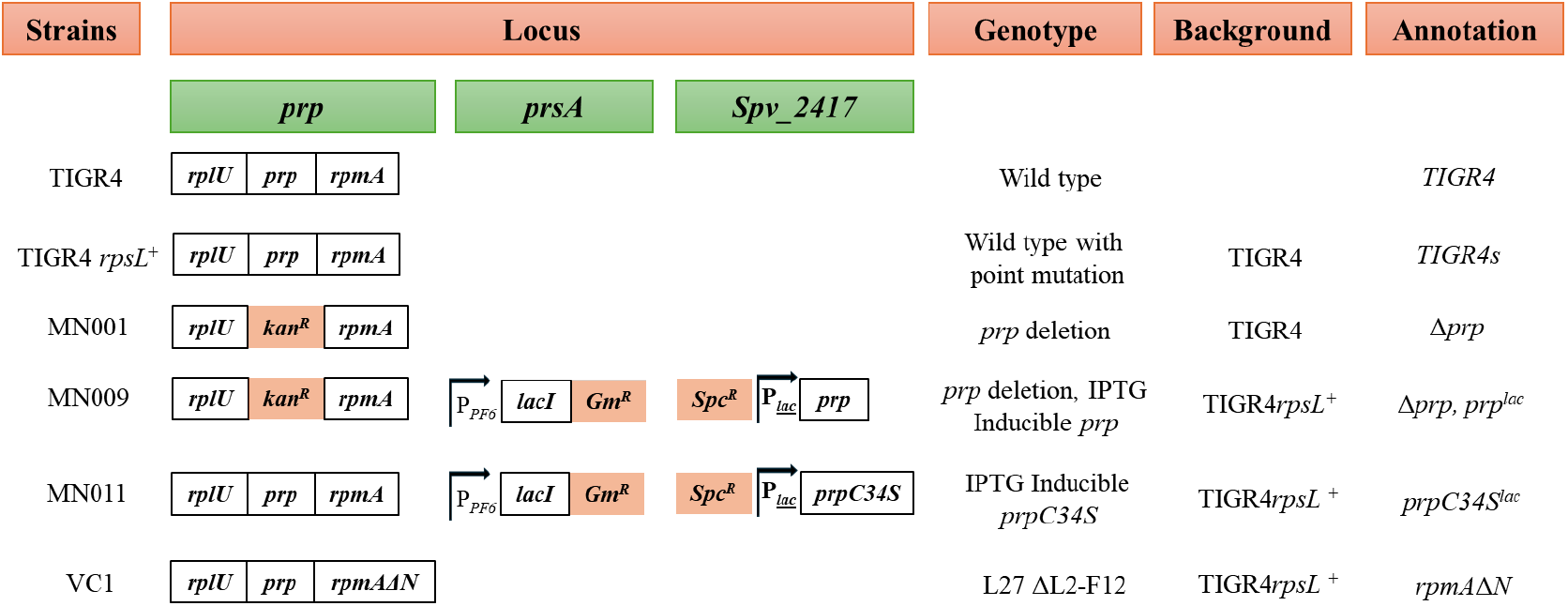
Schematic representation of strains used in this study. Mutations and deletions made in the rplU-prp-rpmA operon, the prsA locus and the Spv_2417 locus are shown. The P_*PF6*_ and P_*lac*_ promoters are shown as square arrows (|^→^), and the antibiotic resistance genes *aph(3’)-IIIa, aac(3)-Ia* and *ant(9)-Ia* are indicated as Kan^R^, Gm^R^ and Spc^R^, respectively.

**Figure 4.**
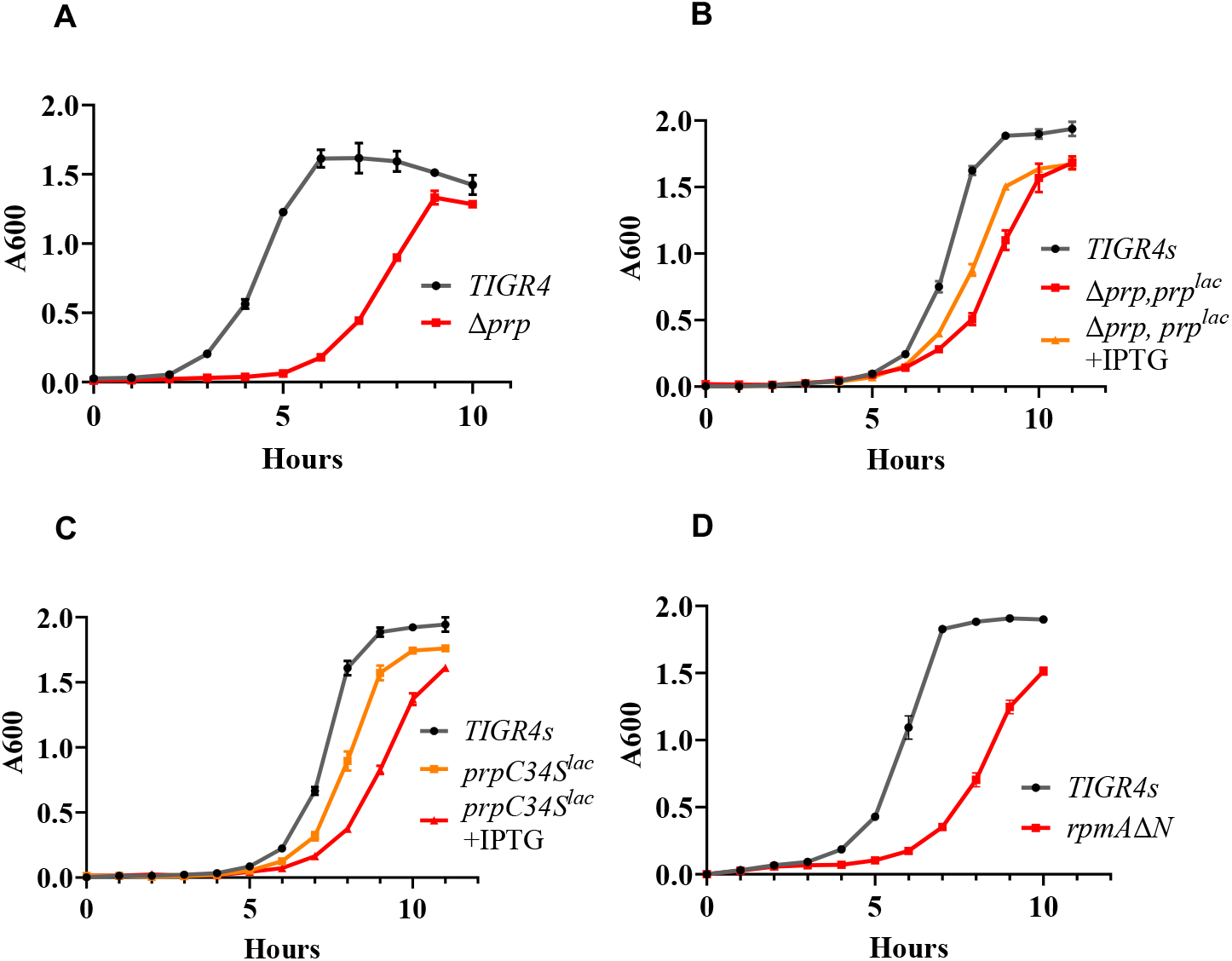
Comparative growth analysis of *Streptococcus pneumoniae* strains. (**A**) Growth curves for the Δ*prp* mutant (strain MN001; red curve) compared to the wildtype strain (TIGR4; black curve). (**B**) Growth curves for the Δ*prp* mutant complemented with an IPTG-inducible *prp* gene inserted at a different locus (MN009) in the presence (orange curve) and absence (red curve) of IPTG compared to the wildtype strain TIGR4s (black curve). (**C**) Growth curves for a strain carrying an IPTG-inducible mutant *prp* gene encoding the protease-defective mutant protein PrpC34S (MN011) in the presence (red curve) and absence (orange curve) of IPTG, compared to the wildtype strain TIGR4s (black curve). (**D**) Growth curves for the N-terminally truncated L27 mutant *rpmA*ΔN (strain VC1; red curve) compared to the wildtype strain (TIGR4s; black curve). Each curve is an average of two experiments, with standard errors shown.

**Table 2.** Growth parameters for *S. pneumoniae* strains.

| Strain | Genotype | Lag phase (hrs) | Increase (-fold)* | Doubling time (min) | Increase (-fold) |
| --- | --- | --- | --- | --- | --- |
| TIGR4 | WT | 3 | 1.0 | 61.0±2.0 | 1 |
| MN001 | $\Delta prp$ | 6 | 2.0 | 62.5±2.5 | 1.03 |
| TIGR4S | *rpsL*+ | 6 | 1.0 | 48.0±3.7 | 1 |
| MN009 – IPTG | $\Delta prp$, *prp*<sup>lac</sup> | 6 | 1.0 | 70.0±1.0 | 1.52 |
| MN009 + IPTG | $\Delta prp$, *prp*<sup>lac</sup> | 6 | 1.0 | 55.6±3.0 | 1.2 |
| MN011 – IPTG | *prpC34S*<sup>lac</sup> | 6 | 1.0 | 50.0±2.0 | 1.17 |
| MN011 + IPTG | *prpC34S*<sup>lac</sup> | 7 | 1.2 | 59.0±2.7 | 1.4 |
| VC1 | *rpmA*ΔN | 7 | 1.2 | 82.0±4.5 | 1.5 |

\* MN001 is relative to TIGR4; other strains are relative to TIGR4S.

To confirm that the growth defect was due to the deletion of *prp*, we made a *prp* complementation strain with an IPTG-inducible *prp* gene. To accomplish this, we first used the plasmid pPEPZ-Plac [27] to introduce a chromosomal copy of *prp* expressing Prp with a C-terminal myc-His_6_ tag under control of the IPTG-inducible *lac* promoter (*P*_lac_) into the *spv_2417* locus in strain TIGR4s. TIGR4s has a point mutation (*rpsL*^+^) in ribosomal protein S12, conferring resistance to streptomycin [28]. Then, the native *prp* allele was replaced with *aphIII* (Kan^R^) in the presence of IPTG, using the same plasmid that was used to generate MN001. Finally, a gene expressing LacI from a constitutive PF6 promoter was introduced into the *prsA* locus using the plasmid pPEPY-PF6-lacI [27]. The resulting strain (MN009; Table 1, Fig. 3), will be referred to as Δ*prp,prp*^lac^.

In the absence of IPTG, the Δ*prp,prp*^lac^ strain showed impaired growth compared to the TIGR4s background strain (Fig. 4B), with a 1.5-fold increase in generation time. The lag phase was not significantly longer, perhaps due to leaky expression of Prp from the *lac* promoter. Upon induction with 5 mM IPTG, growth recovered, albeit not to the same level as the wildtype (Fig. 4B), with a 1.2-fold increase in generation time compared to the wildtype (Table 2). The failure to recover to wildtype levels could be due to distance of the *prp* gene from its original locus, low expression level, or the timing of expression.

### 4. Overexpression of mutant Prp C34S is detrimental to S pneumoniae growth

The *in vitro* results showed that the protease-deficient mutant protein PrpC34S remained bound to SUMO-L27 without cleaving (Fig. 2B). Previous results had shown that expression of PrpC34A had a dominant negative phenotype in *S. aureus* [23]. To check whether expression of PrpC34S would have an equally dominant negative phenotype in *S. pneumoniae*, we used the pPEPZ-Plac plasmid to generate a new strain, in which the C-terminally myc-His_6_-tagged PrpC34S mutant gene was integrated into the *S. pneumoniae* genome in the *spv_2417* locus under control of the *P*_lac_ promoter (strain MN011; *prp*C34S^lac^). This strain also had the *lacI* gene under the constitutive PF6 promoter, as described above (Table 1, Fig. 3).

In the absence of IPTG, strain MN011 exhibited slightly slower growth compared to the wildtype (Fig. 4C). In both cases, the lag phase lasted for six hours, and the *prpC34S*^*lac*^ mutant had a slight increase in generation time compared to wildtype (Table 2). This effect is likely due to leaky expression of PrpC34S from the *lac* promoter. In the presence of IPTG, the lag phase was extended 1.2-fold, the generation time was increased 1.4-fold, and the mutant never reached the same cell density as the wildtype (Fig 4C; Table 2), which is consistent with the results from *S. aureus* [23]. Considering the *in vitro* results presented above, the inhibition of growth by PrpC34S is most likely due to binding of PrpC34S to L27 without release, leading to a failure to assemble functional ribosomes. Note that this strain also expresses wildtype Prp from its native locus, which is most likely the reason the strain is still viable.

### 5. The L27 N-terminal extension is dispensable

To understand the importance of the N-terminal extension of L27, we generated a strain in which the sequence corresponding to the N-terminal 12 amino acids of L27 was deleted, producing L27ΔN (Table 1, Fig. 3). Previous studies had shown that this modification was lethal in *S. aureus* [24]. This strain was made using the EasyJanus cassette, which allows for removal of the antibiotic marker after gene deletion or replacement in a one-step procedure [29]. Surprisingly, the resulting strain (VC1; *rpmA*ΔN; Table 1) was viable, albeit with impaired growth (Fig. 4D), having a longer lag phase and a loner generation time (Table 2). This result suggests that while the L27 N-terminus is important, it is dispensable in *S. pneumoniae*, unlike previous results from *S. aureus* [24].

### 6. Ribosome assembly is normal in the Δprp mutant

In light of the observations in *S. aureus* [23], we hypothesized that the *S. pneumoniae* Δ*prp* mutant would be defective in ribosome assembly. We isolated ribosomes from the TIGR4 wildtype strain and the Δ*prp* mutant strain MN001 at mid-log phase (A_600_=0.6 OD), according to established procedures [30, 31]. The ribosomes were separated on 5–40% sucrose gradients in 0.3 M NH_4_Cl in the presence of 10 mM Mg^2+^, where 70S ribosomes are stable. The sucrose gradient profile was similar in both cases, showing the expected peaks corresponding to 50S subunits and 70S ribosomes (Fig. 5A). The Δ*prp* strain appeared to have a slightly increased amount of 50S subunits compared to the wildtype, but unlike *S. aureus* we did not observe an accumulation of 40S particles. The purified ribosomes looked normal by electron microscopy (Fig. 5B) and displayed a similar banding pattern by SDS-PAGE (Fig. 5C).

**Figure 5.**
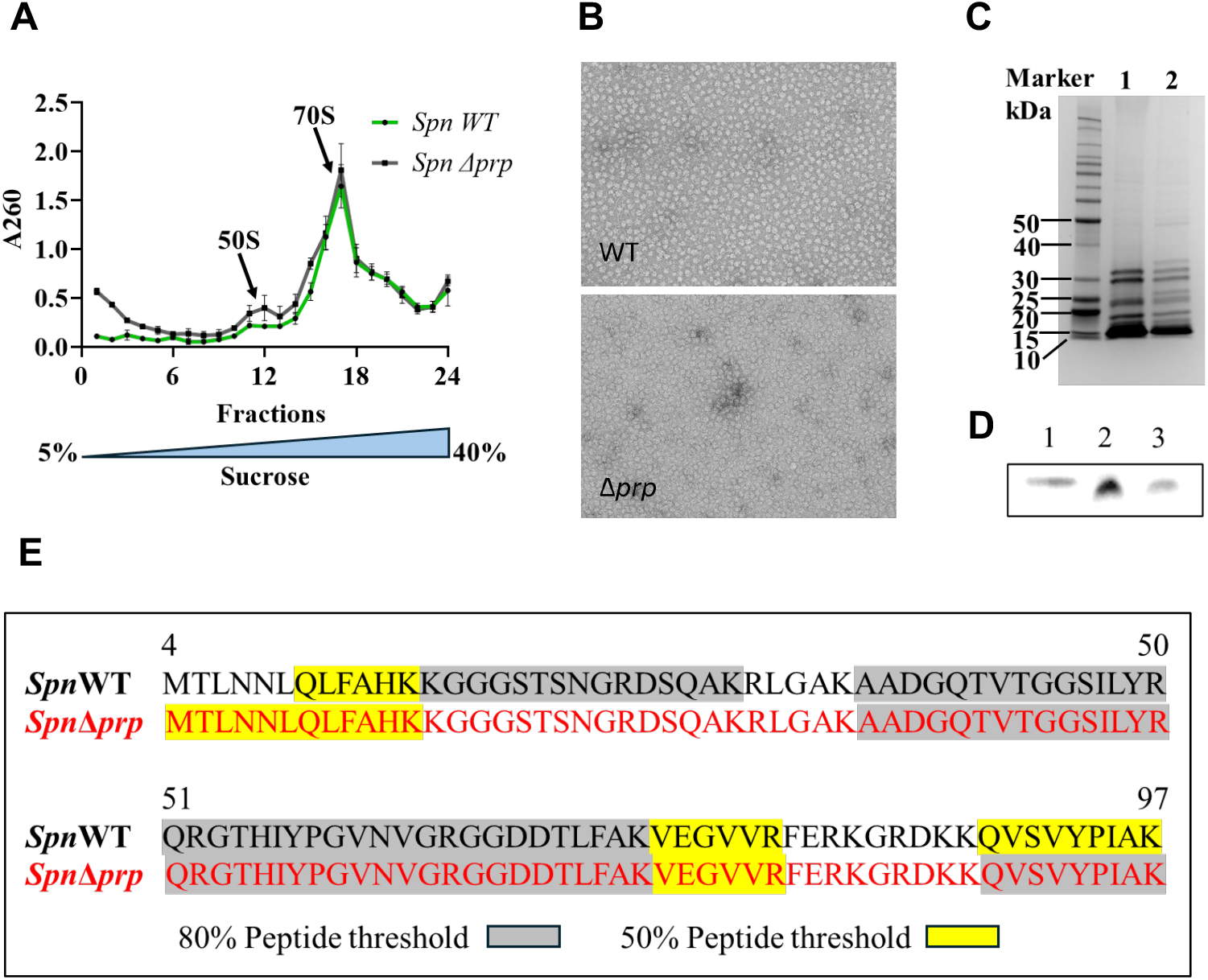
Purification of ribosomes. **(A)** Fractionated sucrose gradient profile for ribosomes purified from wildtype TIGR4 (green curve) and Δ*prp* (MN001; black curve) *S pneumoniae*. **(B)** Negative stain EM of purified ribosomes from TIGR4 (WT) and Δ*prp*. **(C)** SDS-PAGE analysis of purified ribosomes from wildtype (TIGR4; lane 1) and Δ*prp* (MN001; lane 2) strains. **(D)** Western blot analysis using an anti-*S. aureus* L27 antibody. Lane 1, L27ΔN from SUMO-L27 treated with purified Prp; Lane 2, purified ribosomes from wildtype *S. pneumoniae*; Lane 3, purified ribosomes from the Δ*prp* strain. **(E)** Mass spectrometry analysis of L27 in wildtype and Δ*prp* ribosomes. Peptides detected above 80% confidence level are highlighted in gray. Additional peptides detected when the confidence level was lowered to 50% are highlighted in yellow.

To check for the presence or absence of uncleaved L27 in the ribosomes, we separated purified wildtype and MN001 ribosomes by SDS-PAGE and did a Western blot with a polyclonal anti-L27 antibody that we originally raised against *S. aureus* L27 [23], but that also works for *S. pneumoniae* (Fig. 5D). SUMO-L27 cleaved with Prp was used as a control. Surprisingly, there was no difference in the mobility of L27 between the wildtype and Δ*prp* ribosomes, suggesting that the L27 protein had been cleaved even in the absence of Prp.

When the band corresponding to L27 was excised and analyzed by mass spectrometry (MS), no peptides corresponding to the N-terminal extension of L27 were detected at a confidence level >80% (Fig. 5E, Table S1). When the confidence level was lowered to 50%, one peptide corresponding to the N-terminal extension was detected in the Δ*prp* ribosomes, probably corresponding to a background of uncleaved L27 that was not incorporated into ribosomes (Fig. 5E).

### 7. L27 is cleaved in S. pneumoniae cell extracts

The results above suggested that L27 could be cleaved by other proteases present in *S. pneumoniae* in the absence of Prp. To further examine this phenomenon, we mixed the purified SUMO-L27 with lysates from wildtype *S. pneumoniae* (strain TIGR4 and distal lineage strain R6) cells, as well as the MN001 Δ*prp* mutant. As controls, we used a lysate from the Prp-expressing *E. coli* strain described above. The mixtures were incubated for 3 hrs at 37 °C, separated by SDS-PAGE, and probed by Western blotting using either an anti-His_6_ antibody or the anti-L27 antibody mentioned above.

The SUMO-L27 fusion protein was completely cleaved by *E. coli* expressing Prp (Fig. 6A, B). In the presence of the wildtype TIGR4 and D39 lysates, SUMO-L27 was about 50% cleaved after 3 hrs of incubation (Fig. 6A, B). Although the SUMO-L27(1-12) fragment was not detectable after cleavage by the anti-His_6_ antibody (Fig. 6A, S5), presumably due to complete degradation by cellular proteases, it could be observed as a weakening of the SUMO-L27 band. However, a weak band corresponding to the cleaved L27 could be detected when probed with the anti-L27 antibody (Fig. 6B, S5). Importantly, there was no difference in cleavage between the wildtype and Δ*prp* lysates, demonstrating that another protease present in *S. pneumoniae* is able to cleave L27 under these conditions. No cleavage of SUMO-L27 was seen in an *E. coli* lysate in the absence of Prp expression (now shown).

**Figure 6.**
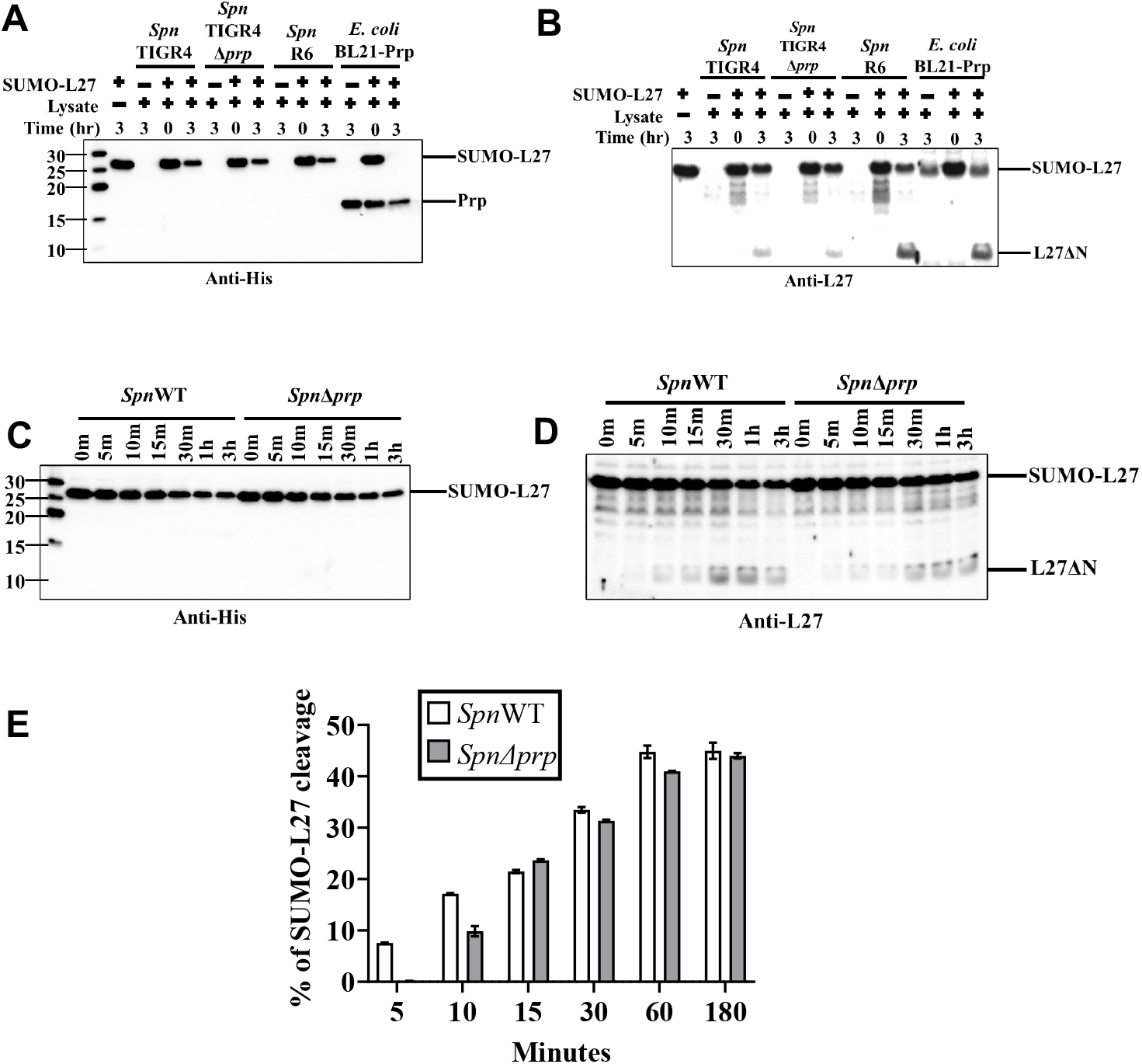
Cleavage of L27 by cell extracts. **(A**,**B)** SUMO-L27 was incubated with lysates of wildtype TIGR4, the TIGR4 Δ*prp* mutant, or *S. pneumoniae* strain R6 for 3 hours at 37 °C. As a positive control for SUMO-L27 cleavage, *E. coli* BL21 lysate expressing hexahistidine-tagged Prp was used. Samples were separated by SDS-PAGE and analyzed by immunoblotting with anti-His (A) and anti-Sau-L27 antibodies (B). **(C**,**D)** A kinetic assay of SUMO-L27 cleavage by cell lysate was performed using the same protocol, followed by immunoblotting with anti-His (C) and anti-S. aureus L27 antibodies (D). **(E)** Amount of SUMO-L27 cleavage, measured as reduction in SUMO-L27 signal detected by the anti-His antibody in (C) from the wildtype (white) and Δ*prp* mutant (gray), quantified by densitometric analysis using ImageJ and plotted over time.

We also carried out an experiment in which SUMO-L27 was mixed with wildtype and Δ*prp* lysates for up to 3 hrs (Fig. 6C,D, S6). This result showed that SUMO-L27 was cleaved efficiently by the Δ*prp* lysate, but with a delayed onset compared to the wildtype (Fig. 6E).

## DISCUSSION

Here, we have examined the cleavage of *S. pneumoniae* ribosomal protein L27 by the Prp protease *in vitro* and *in vivo*. As L27 is a component of the ribosome, *rpmA* is generally considered an essential gene, as shown by transposon mutagenesis in *S. agalactiae* [32] and *S. aureus* [33, 34] and by anti-sense RNA expression in *S. aureus* [35]. In *E. coli*, a Δ*rpmA* mutant was viable, but exhibited four orders of magnitude impaired growth, accumulation of 40S ribosomal large subunit precursors, and ten times reduced peptidyl transferase activity [19, 36]. Transposon mutagenesis and anti-sense RNA expression studies also recorded *prp* as an essential gene in *S. aureus* [34, 35]. In *S. agalactiae, prp* (SAK_1402) was just below the cutoff chosen for essentiality by transposon mutagenesis [32]. In *S. pyogenes*, both *prp* and *rpmA* were assigned “non-conclusive” by transposon mutagenesis [37].

The location of the L27 N-terminus in the peptidyl transferase center is consistent with a strong effect on translation if cleavage is blocked. Using a dual expression system where the wildtype *rpmA* gene could be turned on and off, Wall et al [24] found that *S. aureus* cells expressing either the truncated L27ΔN or an uncleavable form of L27 (L27F12A) were non-viable in the absence of expression of the wildtype allele. The authors of the study were also unable to make a Δ*prp* mutant in *S. aureus*, presumably because it was lethal (unpublished data).

In contrast, a *S. pneumoniae* Δ*prp* mutant was viable. This mutant exhibited delayed growth, but once growth started, it grew with a similar generation time to the wildtype (Fig. 4, Table 2). Moreover, ribosomes formed in the Δ*prp* mutant contained cleaved L27 (Fig. 5), suggesting that another protease was able to carry out this cleavage in the absence of Prp (Fig. 7). This alternate protease is likely less efficient than Prp, resulting in the extended lag phase. This result was further supported by the observed cleavage of the SUMO-L27 fusion protein in a *S. pneumoniae* Δ*prp* lysate, albeit kinetically slower than in the wildtype (Fig. 6E). The *prp* gene is immediately upstream of *rpmA*, which may allow newly synthesized Prp immediate access to L27 for cleavage. Other proteases, encoded at more distant loci, may require more time to locate and cleave L27, thereby leading to the observed delay. Cleavage of L27 appears to be an early event that is a prerequisite for ribosome assembly. Species where *prp* is essential, such as *S. aureus*, might lack an alternate protease capable of cleaving L27 (Fig. 7).

**Figure 7.**
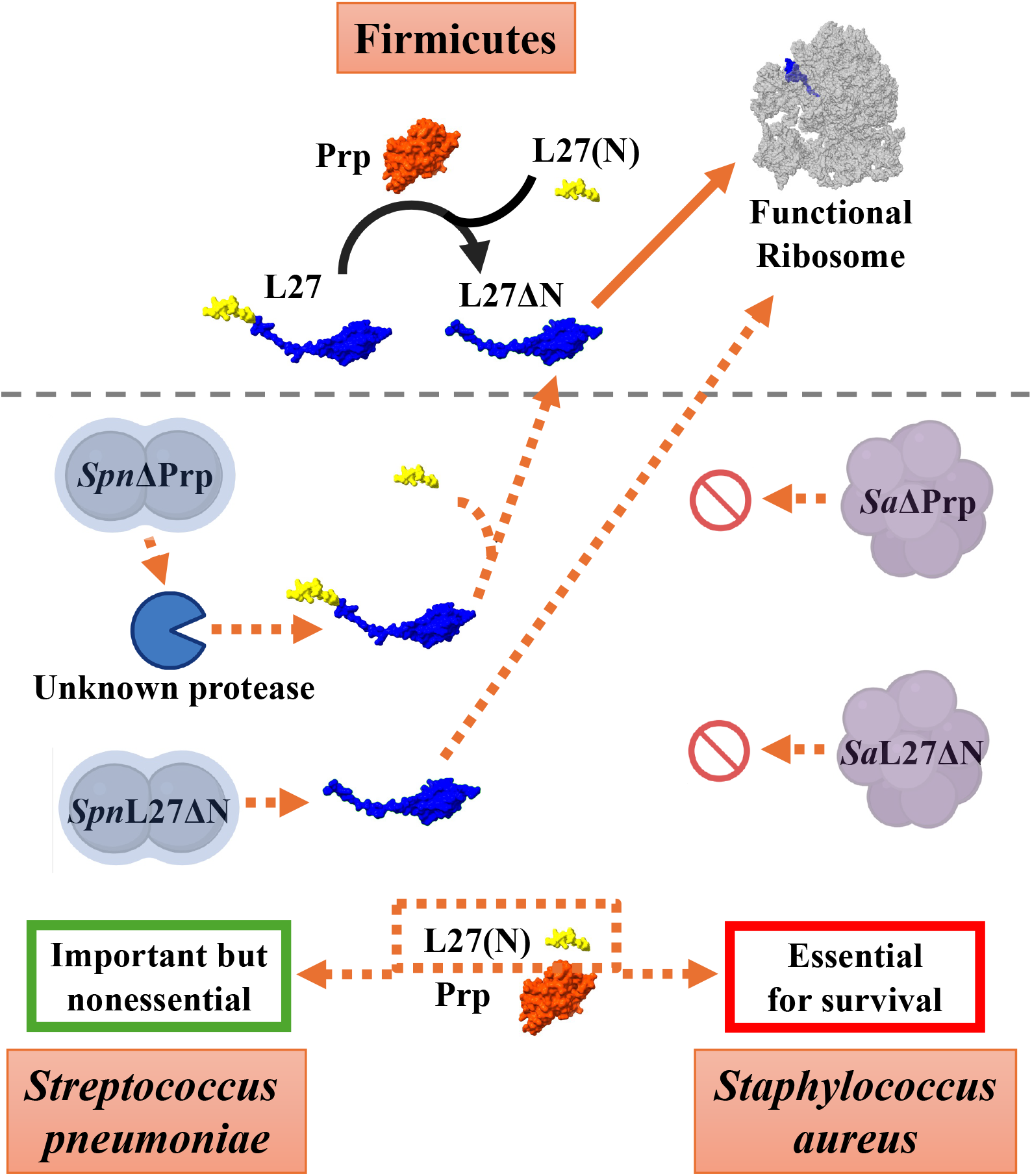
Model for Prp function and L27 cleavage. L27 is normally cleaved by Prp, and only cleaved L27 (L27ΔN) can be incorporated into functional ribosomes. In *S. pneumoniae*, an unknown protease can cleave L27 in the absence of Prp (*Spn*Δprp), and truncated L27 (*Spn*L27ΔN) can also be incorporated. In *S. aureus*, both *Sa*Δprp and *Sa*L27ΔN are deleterious.

Previous studies had shown that expression of the protease-inactive mutant protein PrpC34A or PrpC34S from a plasmid was detrimental to cell growth in *S. aureus*, even in the presence of the wildtype *prp* allele [23]. This strain displayed a distinct ribosome profile, in which 40S ribosomal particles accumulated, presumably corresponding to dead-end intermediates, leading to the hypothesis that L27 cleavage by Prp controlled some step in the ribosome assembly pathway. Similarly, a *S. pneumoniae* strain overexpressing PrpC34S was also impaired in growth. This dominant negative phenotype is most likely due to tight binding of PrpC34S to L27 without cleavage (Fig. 2B). However, the presence of both the wildtype *prp* allele and an alternate protease may be sufficient to prevent aberrant ribosome formation upon PrpC34S overexpression in *S. pneumoniae*.

More surprising was the viability of the L27ΔN mutant, which indicates that the N-terminal extension is not essential in *S. pneumoniae*, unlike in *S. aureus* (Fig. 4D). However, this mutant exhibited slow growth, suggesting that incorporation of the truncated protein into ribosomes was inefficient. In contrast, we were unable to generate a strain in which L27 was mutated (L27F12A) to prevent cleavage by Prp, supporting the idea that cleavage of L27 is essential, whether by Prp or by some other protease (Fig. 7).

We also examined the cleavage of L27 by Prp *in vitro*. Unlike previous studies, which used L27-derived peptides to study the cleavage reaction, we utilized full-length SUMO-tagged L27 and were able to purify a complex between the PrpC34S mutant and SUMO-L27 using SEC (Fig. 2B), which may serve as a valuable tool for future drug design targeting either component. The wildtype Prp protein cleaves rapidly and does not remain associated with L27 after cleavage (Fig. 2A), supporting the notion that Prp does not directly interact with the ribosome.

Although the main role of Prp appears to be the cleavage of L27, it is unclear if it also works on other substrates. Chirgadze et al. [26] suggested that *S. aureus* Prp might be binding to L21, which is encoded by *rplU* in the same operon as *prp* and *rpmA*, but we have no evidence to support this conjecture at present. Recently, Treerat et al [38] described a mutation in the *S. mutans prp* gene (SMU_848) that affected biofilm formation and glucan utilization, possibly through direct cleavage of the glucosyl transferase (Gtc) proteins GtcB and GtcC, which contain a sequence similar to the L27 N-terminal sequence. Although they did not characterize the role of *prp* in ribosome assembly or function in *S. mutans*, the lack of an effect on growth suggests that this species also encodes another protease capable of cleaving L27 [38]. In *S. pneumoniae* TIGR4, we did not identify any convincing matches with sequence homology to the L27 N-terminal sequence, and it is thus unknown if Prp cleaves other substrates.

Our study has shed light on the cleavage of ribosomal protein L27 by Prp which is a requirement for the formation of functional ribosomes. The extended form of L27 and the Prp protease are present in all Firmicutes and many other related bacteria [23], suggesting that this is an evolutionarily conserved and functionally important step. Our results suggest that, in *S. pneumoniae*, only cleaved L27 becomes incorporated into ribosomes, and that the cleavage happens early in the assembly pathway, probably prior to or during formation of the 50S subunit. This differs from the previous study in *S. aureus*, where Prp and the cleavage of L27 appeared to be involved in ribosome assembly [23].

Why these bacteria have the N-terminal extension on L27 in the first place is still a mystery. In *S. aureus*, the presence of an intact N-terminus appeared to be essential [24], but in *S. pneumoniae*, a mutant making only the truncated version of L27 was still viable (Fig. 7). It is possible that the N-terminal extension of L27 plays a role in regulation of ribosomal activity, e.g. under conditions of starvation or other stressors, and may offer protection against certain antibiotics that target the ribosome. While there were no convincing matches to the extended sequences in other proteins in *S. pneumoniae*, it is found in various contexts in other species, including *S. aureus* and *S. mutans*. The presence of the extension in some staphylococcal bacteriophages [23] is especially interesting, suggesting that the sequence is related to an ancient defense mechanism against phages or other environmental stressors. Although not present in strain TIGR4 [39], prophages are common in *S. pneumoniae* isolates [40].

Our results underscore the potential of targeting proteins involved in ribosomal processing and assembly as a strategy to combat antibiotic resistance in *S. pneumoniae*. Prp has been proposed as a target for novel antibiotics [41]. While this approach certainly has merit, it will also be important to understand the implications of L27 cleavage or lack thereof on bacterial dormancy, persistence and antibiotic resistance. While impairing Prp function impacts ribosome activity and downstream cellular processes, it is not lethal in *S. pneumoniae*, highlighting the importance of understanding species-specific differences of therapeutic targets. The present study provides valuable insights for developing novel therapeutic approaches to enhance the effectiveness of current antibiotics and overcome resistance-related challenges.

## ACKNOWLEDGEMENTS

We are grateful to Dr. Jessica Scoffield at UAB for providing strains and plasmids, and for assistance with genetics and bacteriology methods. We appreciate the assistance of Drs. Kyoko Kojima and Jim Mobley in the UAB Proteomics core for mass spectrometry services, and Dr. Peter Prevelige for help with full-length MS. Electron microscopy was done at the UAB Cryo-EM Facility (CEMF), supported by the Institutional Research Core Program, The National Institutes of Health (NIH) grant S10 OD024978 to T.D. and the O’Neal Comprehensive Cancer Center (NIH grant P30 CA013148). This work was funded by an AMC21 pilot grant from the UAB Heersink School of Medicine and NIH grant R21 AI187107 to T.D., and by NIH grant AI114800 to C.J.O.

## MATERIALS AND METHODS

### 1. *Cloning and Site-Directed Mutagenesis in* E. coli

*S. pneumoniae* strain TIGR4 (GenBank ID: AE005672)[39] gene *prp* (SP_1106) was cloned into the pET21a vector using In-Fusion cloning (Takara), incorporating an N-terminal T7 leader sequence followed by the *prp* coding region, a myc tag, a thrombin cleavage site, and a C-terminal hexahistidine (6×His) tag. Similarly, the *S. pneumoniae* TIGR4 gene *rpmA* (SP_1107) was cloned into the pET21a vector preceded by an N-terminal hexahistidine tag and a SUMO domain, also using In-Fusion cloning. Site-directed mutagenesis of both *prp* and *rpmA* was performed using the In-Fusion cloning method (Takara).

### 2. Protein Expression and Purification

Histidine-tagged recombinant proteins were expressed in *E. coli* BL21(DE3) cells following induction with 1 mM IPTG. The expressed proteins were purified using HisPur™ Cobalt Resin (Thermo Scientific). Affinity-purified proteins were concentrated using Amicon centrifugal filter units and further purified by size-exclusion chromatography using Superdex™ 75 Increase columns in a Bio-Rad NGC chromatography system. Eluted protein fractions were collected and concentrated using Amicon filters.

### 3. Enzyme Kinetics and SDS-PAGE Analysis

Prp was serially diluted in assay buffer (25 mM HEPES, pH 7.4; 0.3 M NaCl) and incubated with increasing concentrations of SUMO-L27 at 37 °C either for 1 hour or for various time intervals. Reactions were stopped by adding Laemmli sample buffer followed by boiling at 95 °C for 4 minutes. Reaction products were analyzed by SDS-PAGE, and band intensities were quantified by densitometric analysis using ImageJ. To determine the Michaelis constant (Km), 1 µM Prp was incubated with varying concentrations of SUMO-L27 for 15 minutes at 37 °C. The reactions were terminated as described above, and product formation was analyzed by SDS-PAGE followed by densitometric quantification using ImageJ.

### 4. Binding assay and Size exclusion Chromatography

Equimolar amounts of SUMO-L27 and Prp were mixed in assay buffer (25 mM HEPES, pH 7.4; 0.3 M NaCl) and incubated at 37 °C for 2 hours with gentle rotation. The reaction mixture was then applied to a Superdex™ 75 Increase size-exclusion chromatography column (GE Healthcare) pre-equilibrated with assay buffer. Eluted fractions were collected and analyzed by SDS-PAGE.

### 5. Generation of S. pneumoniae mutants

I. Deletion of *prp*: An in-frame deletion of the *prp* gene was performed in *S. pneumoniae* TIGR4 using the pPEPZ-Plac plasmid [27]. A 2,873 bp kanamycin resistance cassette was constructed by assembling a 1010 bp left homology arm (upstream of prp), a 1068 bp right homology arm (downstream of prp), and a 795 bp kanamycin resistance gene (*aphIII)*. These components were joined via In-Fusion PCR and cloned into the pPEPZ-Plac plasmid. The recombinant plasmid was transformed into *S. pneumoniae* TIGR4, and transformants were selected on blood agar plates containing kanamycin, yielding the Δ*prp* strain MN001 (Table 1).
II. Introduction of wildtype and mutant *prp* alleles: Wildtype and mutant alleles of *prp* were introduced into *S. pneumoniae* serotype 4, strain TIGR4 *rpsL*+, which harbors a point mutation in gene *rpsL* (protein S12), conferring resistance to streptomycin [28]. The wildtype *prp* gene was cloned into the pPEPZ-Plac plasmid [27] under the control of the *lac* promoter and containing a spectinomycin resistance marker. This construct integrates into the non-essential pseudogene *spv_2417*. The *prp* gene was then replaced with the kanamycin resistance gene, as described above. Finally, the plasmid pPEPY-PF6-lacI [27], encoding the *lacI* gene under gentamicin selection, was integrated downstream of the *prs1* locus to regulate *lac* promoter activity, resulting in strain MN009 (Table 1). Similarly, a mutant *prp* gene expressing PrpC34S protein under control of the *lac* promoter was introduced into the *spv_2417* locus using pPEPZ-Plac, followed by introduction of *lacI* using pPEPY-PF6-lacI, yielding strain MN011 (Table 1). Both MN009 and MN011 constructs included a C-terminal myc tag, a thrombin cleavage site, and a 6×His tag after the *prp* coding sequence.
III. N-terminal truncation of L27: The strain VC1, containing a truncated version of the *rpmA* gene in its native locus, was generated in the TIGR4 *rpsL*^+^ using the easyJanus method [29]. This strain produces a truncated version of ribosomal protein L27 lacking residues 2–12. To generate this strain, synthetic DNA fragments were obtained from Twist Bioscience (South San Francisco, CA, USA). DNA assembly was performed using the NEBuilder HiFi DNA Assembly Master Mix kit (New England Biolabs). The assembled constructs were introduced into competent cells in a single transformation step, and transformants were selected on kanamycin-containing agar plates, as previously described [29].

### 6. Bacterial growth assay

Growth assays for wild-type and mutant *S. pneumoniae* strains were performed in Todd-Hewitt broth supplemented with 2% yeast extract (THY). Briefly, cells were grown overnight on blood agar plates (Remel) at 37 °C with 5% CO_2_. One-fourth of the plate surface was scraped using a sterile cell scraper and used to inoculate 10 mL of THY as a starter culture. The starter culture was grown until it reached an optical density (OD) of 0.3 at 600 nm wavelength (A_600_). A 2.5% inoculum from the starter culture was used to inoculate 200 mL of fresh THY medium. Bacterial growth was monitored by measuring A_600_ every hour for 10 hours. For IPTG-inducible mutants, cultures were grown either in the absence of IPTG or in the presence of 5 mM IPTG. All experiments were performed in triplicate.

### 7. Ribosome purification

Ribosomes were purified from *S. pneumoniae* wildtype and Δ*prp* strains. Briefly, 500 mL cultures were grown to A_600_ = 0.4 OD and harvested by centrifugation. Cell pellets were resuspended in 15 mL lysis buffer (20 mM Tris-HCl, pH 7.5; 300 mM NH_4_Cl; 15 mM MgCl_2_; 0.5 mM EDTA; 6 mM β-mercaptoethanol) and lysed by three passes through an Avestin Emulsiflex B15 homogenizer. The lysate was clarified by centrifugation at 25,000 × g for 30 minutes at 4 °C. The clarified lysate (15 mL) was carefully layered onto 10 mL of sucrose cushion buffer (1.1 M sucrose in lysis buffer) and centrifuged at 50,000 rpm for 16 hours at 4 °C in a Ti70 rotor. After centrifugation, the supernatant was discarded, and the ribosomal pellet was resuspended in 10 mL of lysis buffer. A 150 µL aliquot of the ribosome preparation was then layered onto a 5–40% (w/v) sucrose gradient prepared in lysis buffer and centrifuged at 75,500 × g for 16 hours at 4 °C in an SW41 rotor. Fractions containing 70S ribosomal particles were identified, pooled, dialyzed against lysis buffer, and concentrated using a 100 kDa molecular weight cutoff membrane. The ribosome samples were negatively stained with 1% (w/v) uranyl acetate and examined by electron microscopy using an FEI Tecnai F20 transmission electron microscope.

### 8. Mass spectrometry

Purified SUMO-L27 and Prp were mixed and incubated at 37 °C for 1 hour to allow cleavage. Following incubation, the release of the N-terminal fragment of L27 was analyzed by mass spectrometry using a Waters-Synapt ESI-TOF-MS instrument. To examine the state of L27 in ribosomes, wildtype and Δ*prp* 70S ribosomes were separated by SDS-PAGE on a 10% Bis-Tris gel and stained with colloidal Coomassie blue. A segment corresponding to the visible bands was extracted, digested with trypsin and analyzed by LC-ESI-MS/MS using a Thermo Orbitrap Velos Pro spectrometer.

### 9. In Vitro Cleavage Assay by cell lysate and Immunoblot Analysis

Cells were harvested at A_600_=0.6 OD by centrifugation. The cell pellets were resuspended in assay buffer (25 mM HEPES, pH 7.4; 0.3 M NaCl) and lysed in an Avestin Emulsiflex B15 homogenizer. The lysate was clarified by centrifugation, and the supernatant was filtered through a 0.2 µm membrane filter. Protein concentration in the lysate was determined using the Bradford assay (Bio-Rad) and normalized to 2 mg/mL. For the cleavage assay, 3.5 µM of purified SUMO-L27 in assay buffer was mixed with the cell lysate and incubated at 37 °C for varying time intervals with gentle rotation. The control sample was boiled instantly with Laemmli loading buffer after mixing the protein and lysate. Samples were analyzed by immunoblotting using either an anti-His primary antibody (Invitrogen, 37-2900) or a custom rabbit anti-*S. aureus* L27 antibody [23]. A horseradish peroxidase (HRP)-conjugated anti-rabbit secondary antibody was used for detection. Signal development was carried out using a chemiluminescent substrate composed of luminol and hydrogen peroxide (Fisher Scientific).

## REFERENCES

1. Kadioglu A., Weiser J.N., Paton J.C. & Andrew P.W. (2008). The role of Streptococcus pneumoniae virulence factors in host respiratory colonization and disease. Nat Rev Microbiol. 6, 288–301.

2. Mitchell A.M. & Mitchell T.J. (2010). Streptococcus pneumoniae: virulence factors and variation. Clin Microbiol Infect. 16, 411–418.

3. Dion C.F. & Ashurst J.V. (2025) tStreptococcus pneumoniae. In StatPearls StatPearls Publishing, Treasure Island (FL).

4. O’Brien, KL, Wolfson, LJ, Watt, JP, Henkle, E, Deloria-Knoll, M, McCall, N, Lee, E, Mulholland, K, Levine, OS, Cherian, T & Hib and Pneumococcal Global Burden of Disease Study Team (2009). Burden of disease caused by Streptococcus pneumoniae in children younger than 5 years: global estimates. Lancet. 374, 893–902.

5. Roos K.L. (2000). Acute bacterial meningitis. Semin Neurol. 20, 293–306.

6. Cherazard R., Epstein M., Doan T.L., Salim T., Bharti S. & Smith M.A. (2017). Antimicrobial Resistant Streptococcus pneumoniae: Prevalence, Mechanisms, and Clinical Implications. Am J Ther. 24, e361–e369.

7. Narciso A.R., Dookie R., Nannapaneni P., Normark S. & Henriques-Normark B. (2025). Streptococcus pneumoniae epidemiology, pathogenesis and control. Nat Rev Microbiol. 23, 256–271.

8. Lui G.C.Y. & Lai C.K.C. (2025). Community acquired pneumonia due to antibiotic resistant-Streptococcus pneumoniae : diagnosis, management and prevention. Curr Opin Pulm Med. 31, 211–217.

9. Willenborg J., Willms D., Bertram R., Goethe R. & Valentin-Weigand P. (2014). Characterization of multi-drug tolerant persister cells in Streptococcus suis. BMC Microbiol. 14, 120.

10. Hernandez-Morfa M., Reinoso-Vizcaíno N.M., Olivero N.B., Zappia V.E., Cortes P.R., Jaime A. & Echenique J. (2022). Host Cell Oxidative Stress Promotes Intracellular Fluoroquinolone Persisters of Streptococcus pneumoniae. Microbiol Spectr. 10, e0436422.

11. Wilson D.N. (2014). Ribosome-targeting antibiotics and mechanisms of bacterial resistance. Nat Rev Microbiol. 12, 35–48.

12. Lin J., Zhou D., Steitz T.A., Polikanov Y.S. & Gagnon M.G. (2018). Ribosome-Targeting Antibiotics: Modes of Action, Mechanisms of Resistance, and Implications for Drug Design. Annu Rev Biochem. 87, 451–478.

13. Auerbach T., Bashan A., Harms J., Schluenzen F., Zarivach R., Bartels H., Agmon I., Kessler M., Pioletti M., Franceschi F. & Yonath A. (2002). Antibiotics targeting ribosomes: crystallographic studies. Curr Drug Targets Infect Disord. 2, 169–186.

14. Yonath A. (2005). Antibiotics targeting ribosomes: resistance, selectivity, synergism and cellular regulation. Annu Rev Biochem. 74, 649–679.

15. Tebano G., Zaghi I., Baldasso F., Calgarini C., Capozzi R., Salvadori C., Cricca M. & Cristini F. (2024). Antibiotic Resistance to Molecules Commonly Prescribed for the Treatment of Antibiotic-Resistant Gram-Positive Pathogens: What Is Relevant for the Clinician. Pathogens. 13, 88.

16. Shajani Z., Sykes M.T. & Williamson J.R. (2011). Assembly of bacterial ribosomes. Annu Rev Biochem. 80, 501–526.

17. Kaczanowska M. & Ryden-Aulin M. (2007). Ribosome biogenesis and the translation process in Escherichia coli. Microbiol Mol Biol Rev. 71, 477–494.

18. Voorhees R.M., Weixlbaumer A., Loakes D., Kelley A.C. & Ramakrishnan V. (2009). Insights into substrate stabilization from snapshots of the peptidyl transferase center of the intact 70S ribosome. Nat Struct Mol Biol. 16, 528–533.

19. Wower I.K., Wower J. & Zimmermann R.A. (1998). Ribosomal protein L27 participates in both 50 S subunit assembly and the peptidyl transferase reaction. J Biol Chem. 273, 19847–19852.

20. Maguire B.A., Beniaminov A.D., Ramu H., Mankin A.S. & Zimmermann R.A. (2005). A protein component at the heart of an RNA machine: the importance of protein L27 for the function of the bacterial ribosome. Mol Cell. 20, 427–435.

21. Maracci C., Wohlgemuth I. & Rodnina M.V. (2015). Activities of the peptidyl transferase center of ribosomes lacking protein L27. RNA. 21, 2047–2052.

22. Poliakov A., Chang J.R., Spilman M.S., Damle P.K., Christie G.E., Mobley J.A. & Dokland T. (2008). Capsid size determination by Staphylococcus aureus pathogenicity island SaPI1 involves specific incorporation of SaPI1 proteins into procapsids. J. Mol. Biol. 380, 465–475.

23. Wall E.A., Caufield J.H., Lyons C.E., Manning K.A., Dokland T. & Christie G.E. (2015). Specific N-terminal cleavage of ribosomal protein L27 in Staphylococcus aureus and related bacteria. Mol Microbiol. 95, 258–269.

24. Wall E.A., Johnson A.L., Peterson D.L. & Christie G.E. (2017). Structural modeling and functional analysis of the essential ribosomal processing protease Prp from Staphylococcus aureus. Mol Microbiol. 104, 520–532.

25. Hotinger J.A., Pendergrass H.A., Peterson D., Wright H.T. & May A.E. (2022). Phage-Related Ribosomal Protease (Prp) of Staphylococcus aureus: In Vitro Michaelis-Menten Kinetics, Screening for Inhibitors, and Crystal Structure of a Covalent Inhibition Product Complex. Biochemistry. 61, 1323–1336.

26. Chirgadze Y.N., Clarke T.E., Romanov V., Kisselman G., Wu-Brown J., Soloveychik M., Chan T.S., Gordon R.D., Battaile K.P., Pai E.F. & Chirgadze N.Y. (2015). The structure of SAV1646 from Staphylococcus aureus belonging to a new ribosome-associated’ subfamily of bacterial proteins. Acta Crystallogr D Biol Crystallogr. 71, 332–337.

27. Keller L.E., Rueff A.S., Kurushima J. & Veening J.W. (2019). Three New Integration Vectors and Fluorescent Proteins for Use in the Opportunistic Human Pathogen. Genes (Basel). 10, 394.

28. Sung C.K., Li H., Claverys J.P. & Morrison D.A. (2001). An rpsL cassette, janus, for gene replacement through negative selection in Streptococcus pneumoniae. Appl Environ Microbiol. 67, 5190–5196.

29. Chembilikandy V., D’Mello A., Tettelin H., Martínez E. & Orihuela C.J. (2024). Streamlining marker-less allelic replacement in Streptococcus pneumoniae through a single transformation step strategy: easyJanus. Appl Environ Microbiol. 90, e0101024.

30. Blaha G., Stelzl U., Spahn C.M., Agrawal R.K., Frank J. & Nierhaus K.H. (2000). Preparation of functional ribosomal complexes and effect of buffer conditions on tRNA positions observed by cryoelectron microscopy. Methods Enzymol. 317, 292–309.

31. Khusainov I., Vicens Q., Bochler A., Grosse F., Myasnikov A., Ménétret J.F., Chicher J., Marzi S., Romby P., Yusupova G., Yusupov M. & Hashem Y. (2016). Structure of the 70S ribosome from human pathogen Staphylococcus aureus. Nucleic Acids Res. 44, 10491–10504.

32. Hooven T.A., Catomeris A.J., Akabas L.H., Randis T.M., Maskell D.J., Peters S.E., Ott S., Santana-Cruz I., Tallon L.J., Tettelin H. & Ratner A.J. (2016). The essential genome of Streptococcus agalactiae. BMC Genomics. 17, 406.

33. Chaudhuri R.R., Allen A.G., Owen P.J., Shalom G., Stone K., Harrison M., Burgis T.A., Lockyer M., Garcia-Lara J., Foster S.J., Pleasance S.J., Peters S.E., Maskell D.J. & Charles I.G. (2009). Comprehensive identification of essential Staphylococcus aureus genes using Transposon-Mediated Differential Hybridisation (TMDH). BMC Genomics. 10, 291.

34. Fey P.D., Endres J.L., Yajjala V.K., Widhelm T.J., Boissy R.J., Bose J.L. & Bayles K.W. (2013). A genetic resource for rapid and comprehensive phenotype screening of nonessential Staphylococcus aureus genes. MBio. 4, e00537–12.

35. Ji Y., Woodnutt G., Rosenberg M. & Burnham M.K. (2002). Identification of essential genes in Staphylococcus aureus using inducible antisense RNA. Methods Enzymol. 358, 123–128.

36. Trobro S. & Aqvist J. (2008). Role of ribosomal protein L27 in peptidyl transfer. Biochemistry. 47, 4898–4906.

37. Le Breton Y., Belew A.T., Valdes K.M., Islam E., Curry P., Tettelin H., Shirtliff M.E., El-Sayed N.M. & McIver K.S. (2015). Essential Genes in the Core Genome of the Human Pathogen Streptococcus pyogenes. Sci Rep. 5, 9838.

38. Treerat P., de Mattos C., Burnside M., Zhang H., Zhu Y., Zou Z., Anderson D., Wu H., Merritt J. & Kreth J. (2024). Ribosomal-processing cysteine protease homolog modulates Streptococcus mutans glucan production and interkingdom interactions. J Bacteriol. 206, e0010424.

39. Tettelin H., Nelson K.E., Paulsen I.T., Eisen J.A., Read T.D., Peterson S., Heidelberg J., DeBoy R.T., Haft D.H., Dodson R.J., Durkin A.S., Gwinn M., Kolonay J.F., Nelson W.C., Peterson J.D., Umayam L.A., White O., Salzberg S.L., Lewis M.R., Radune D., Holtzapple E., Khouri H., Wolf A.M., Utterback T.R., Hansen C.L., McDonald L.A., Feldblyum T.V., Angiuoli S., Dickinson T., Hickey E.K., Holt I.E., Loftus B.J., Yang F., Smith H.O., Venter J.C., Dougherty B.A., Morrison D.A., Hollingshead S.K. & Fraser C.M. (2001). Complete genome sequence of a virulent isolate of Streptococcus pneumoniae. Science. 293, 498–506.

40. Garriss G. & Henriques-Normark B. (2020). Lysogeny in Streptococcus pneumoniae. Microorganisms. 8, 1546.

41. Hotinger J.A., Gallagher A.H. & May A.E. (2022). Phage-Related Ribosomal Proteases (Prps): Discovery, Bioinformatics, and Structural Analysis. Antibiotics (Basel). 11, 1109.

42. Liu X., Gallay C., Kjos M., Domenech A., Slager J., van Kessel S.P., Knoops K., Sorg R.A., Zhang J.R. & Veening J.W. (2017). High-throughput CRISPRi phenotyping identifies new essential genes in Streptococcus pneumoniae. Mol Syst Biol. 13, 931.

