## Supplementary Figures and Table for "Cleavage of *Streptococcus pneumoniae* ribosomal protein L27 by the Prp protease"

**S1**

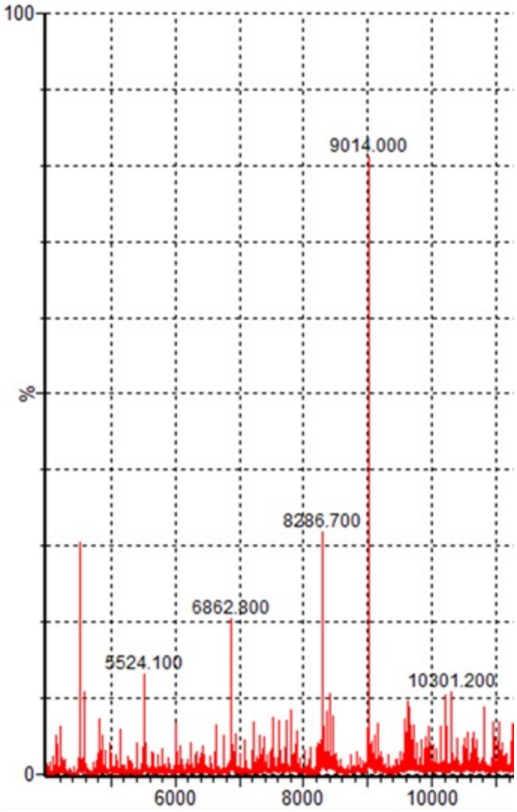


**Figure S1. Mass spectrometry of cleaved SUMO-L27.** Intact proteins analysis by ESI-MS of SUMO-L27 cleaved with Prp, showing the expected 9,014 Da peak corresponding to cleaved L27.

S2

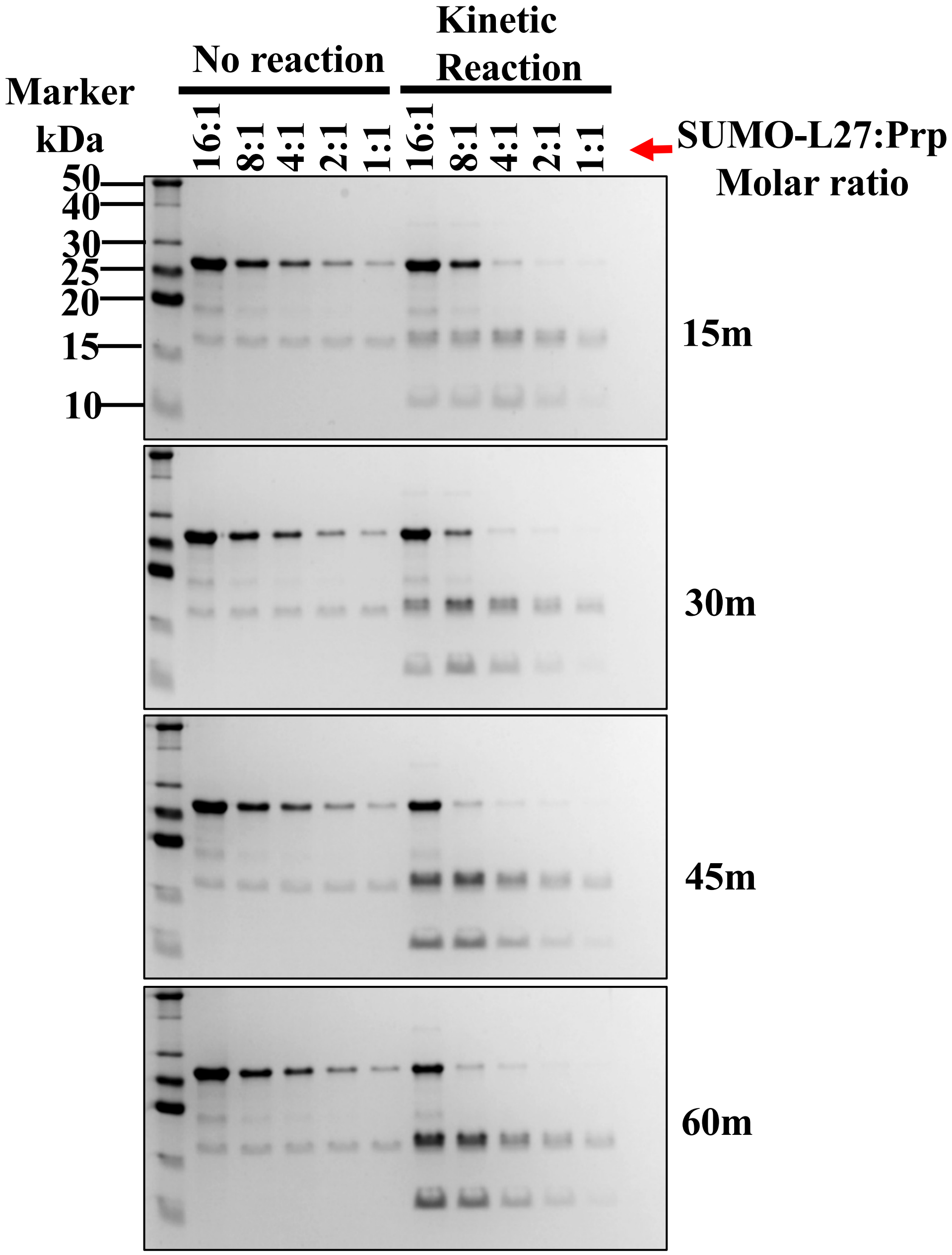


**Figure S2. Time course of SUMO-L27 cleavage in vitro.** Coomassie stained SDS-PAGE separation of SUMO-L27 cleaved with Prp at different molar ratios for 15, 30, 45 and 60 mins (kinetic reaction). Each gel includes a control (no reaction), in which the proteins were mixed directly in sample buffer and boiled so that no cleavage would occur.

S3

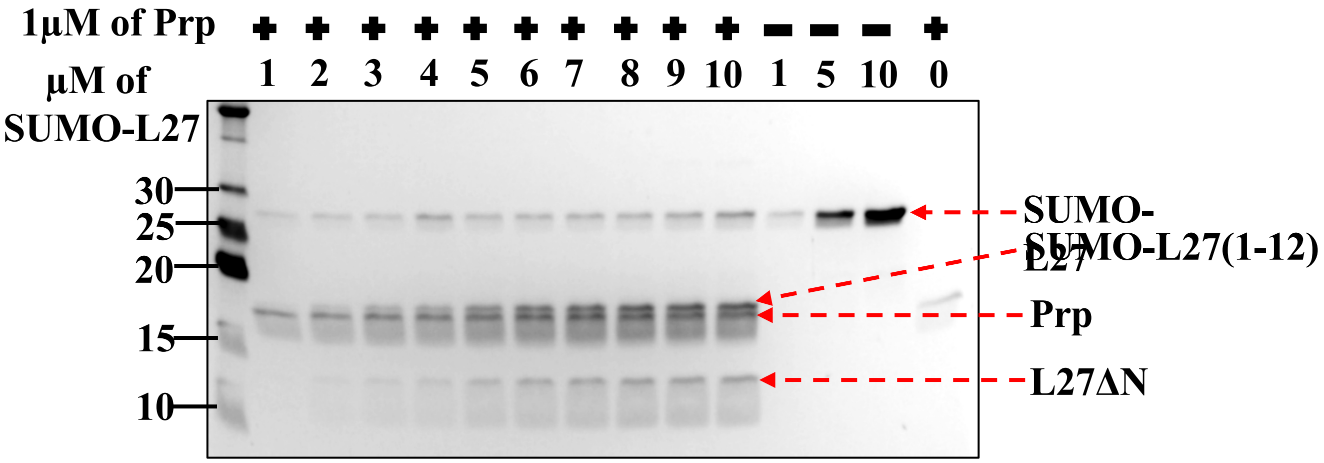


**Figure S3. Concentration dependence of SUMO-L27 cleavage.** Coomassie stained SDS-PAGE separation showing cleavage of varying concentrations of SUMO-L27 from 1 to 10 µM by 1 µM of Prp. Bands corresponding to uncleaved SUMO-L27, the two cleavage products SUMO-L27(1-12) and L27ΔN, and Prp are indicated.

S4A

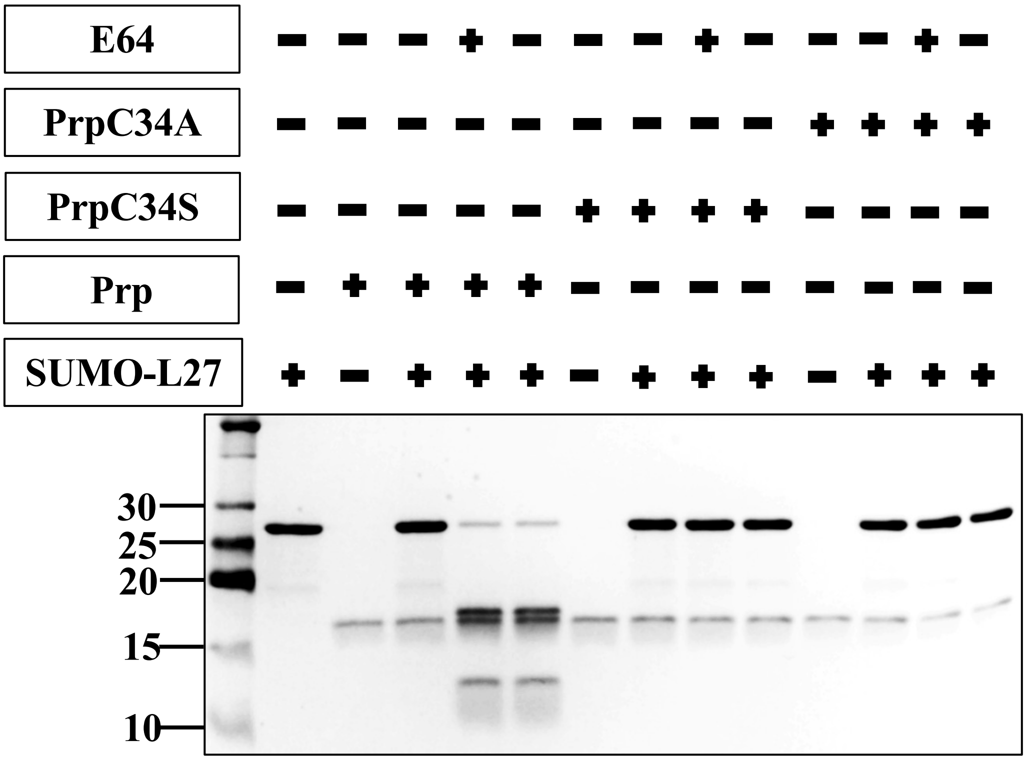
E64 − − − + − − − + − − − + −
PrpC34A − − − − − − − − − + + + +
PrpC34S − − − − − + + + + − − − −
Prp − + + + + − − − − − − − −
SUMO-L27 + − + + + − + + + − + + +
30
25
20
15
10

S4B

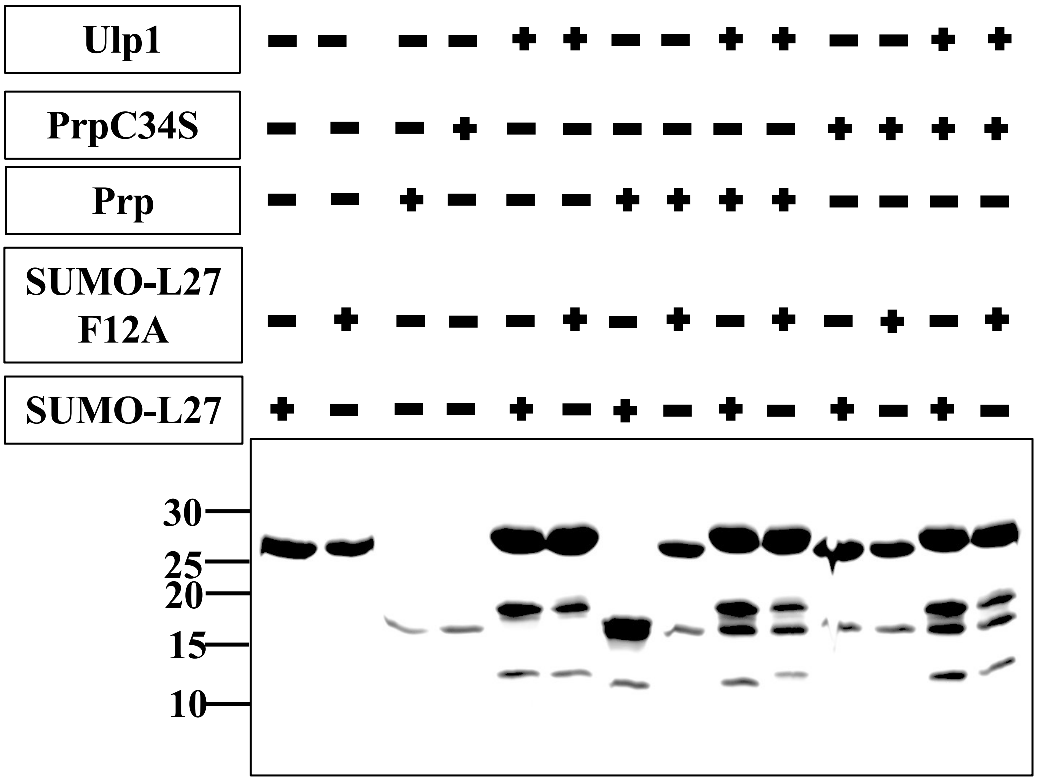
Ulp1 − − − − + + − − + + − − + +
PrpC34S − − − + − − − − − − + + + +
Prp − − + − − − + + + + − − − −
SUMO-L27 F12A − + − − − + − + − + − + − +
SUMO-L27 + − − − + − + − + − + − + −
30
25
20
15
10

**Figure S4. Cleavage of SUMO-L27.** (**A**) Coomassie stained SDS-PAGE separation of 2 μM SUMO-L27 after incubating with 2 μM of either Prp, PrpC34S or PrpC34A for 1 hr at 37 °C. Cysteine protease inhibitor E-64 was added (1 mM) to the reaction in the indicated lanes. (**B**) Coomassie stained SDS-PAGE separation of 10 μM SUMO-L27 or SUMO-L27F12A incubated with 10 μM Prp, PrpC34S or SUMO protease (Ulp1) for 2 hrs at 37 °C.

S5

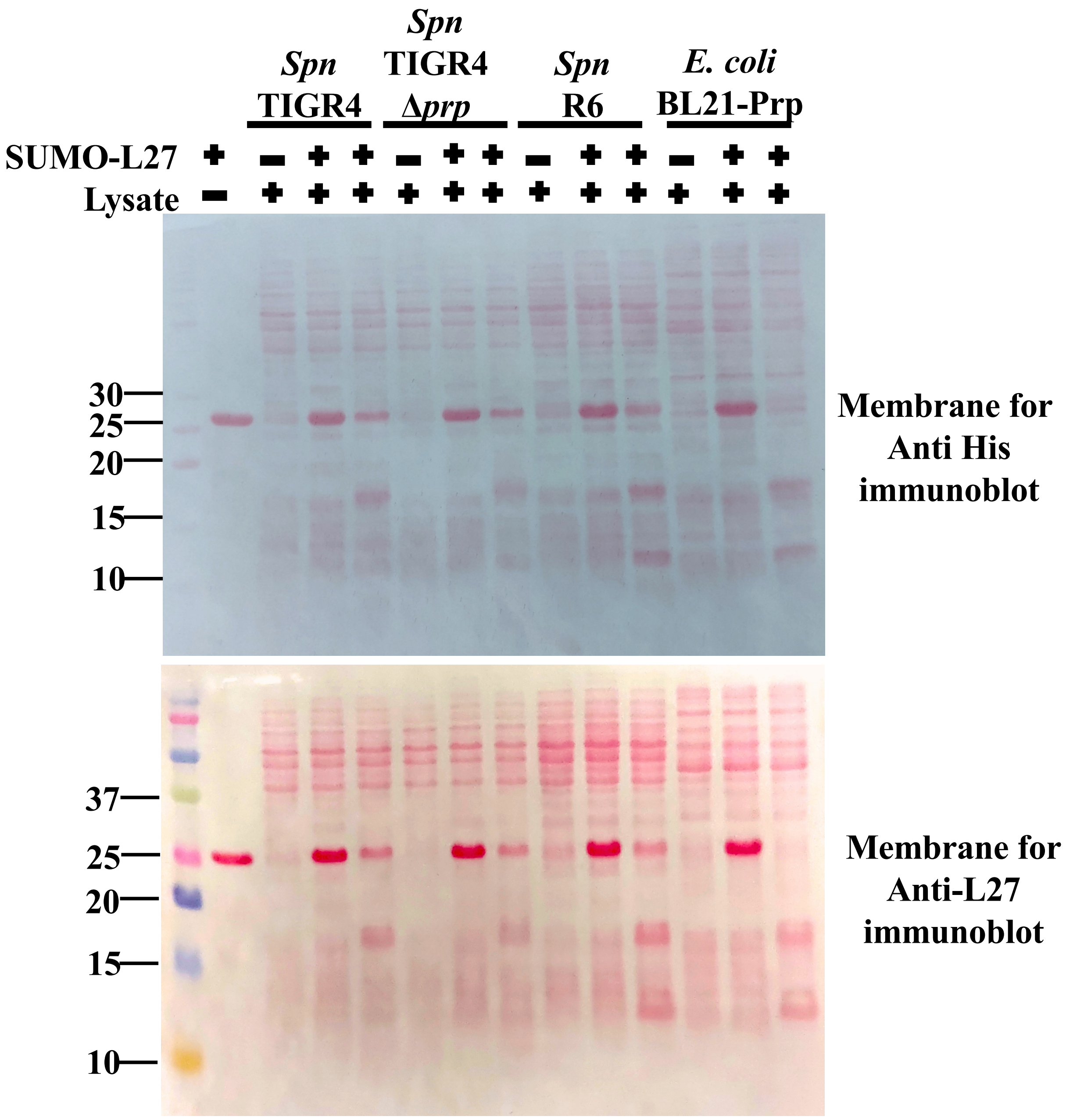


**Figure S5. Cleavage of SUMO-L27 by cell lysates.** Ponceau stained Western blot membranes of SUMO-L27 cleaved by cell lysates of *S. pneumoniae* strains TIGR4, MN001 (Δ*prp*), R6 or *E. coli* expressing Prp. Corresponds to the Western blots in Figure 6A (stained with anti-His antibody) and 6B (anti-L27).

S6

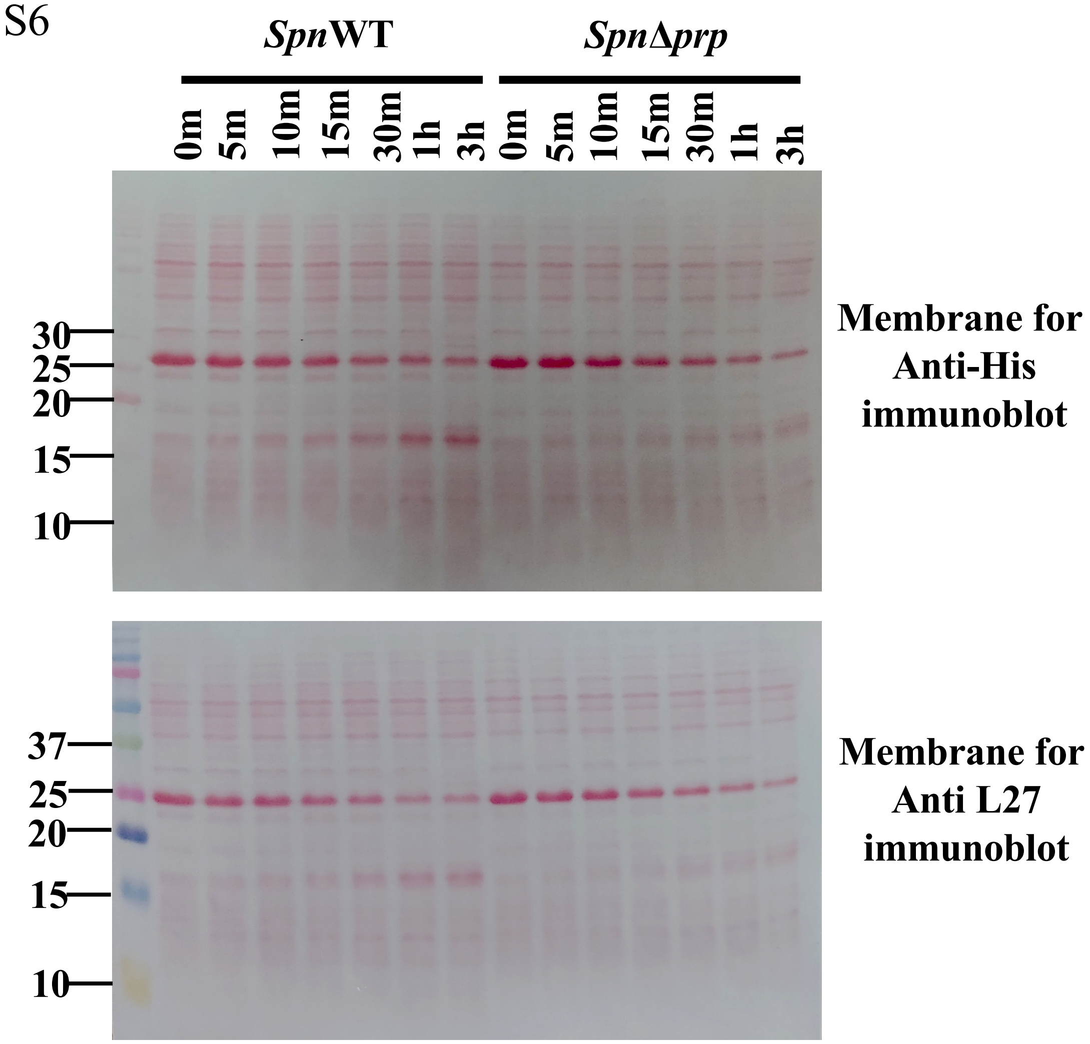


**Figure S6. Cleavage of SUMO-L27 by cell lysates.** Ponceau stained Western blot membranes of SUMO-L27 cleaved by *S. pneumoniae* strains TIGR4 (*Spn*WT) or MN001 (*Spn*Δ*prp*) for up to 3 hrs. Corresponds to the Western blots in Figure 6C (anti-His antibody) and 6D (anti-L27).

**Table S1:** Total spectrum counts of L27 peptides.

| Peptides | Total Spectrum Count | |
|---|---|---|
| | SpnWT | SpnΔprp |
| AADGQTVTGGSILYR | 22 | 28 |
| DDTLFAK | 3 | 5 |
| DTLFAK | 3 | 2 |
| GDDTLFAK | 5 | 3 |
| GGDDTLFAK | 35 | 27 |
| GGGSTSNGRDSQAK | 1 | 0 |
| GQTVTGGSILYR | 0 | 1 |
| GTHIYPGVNVGR | 26 | 27 |
| GVNVGR | 3 | 3 |
| KGGGSTSNGRDSQAK | 1 | 0 |
| MTLNNLQLFAHK | 0 | 1 |
| PGVNVGR | 2 | 1 |
| QLFAHK | 1 | 0 |
| QRGTHIYPGVNVGR | 3 | 1 |
| QVSVYPIAK | 2 | 3 |
| VEGVVR | 31 | 64 |
| VNVGRGGDDTLFAK | 1 | 1 |
